# The late blooming amphipods: global change promoted post-Jurassic ecological radiation despite Palaeozoic origin

**DOI:** 10.1101/675140

**Authors:** Denis Copilaş-Ciocianu, Špela Borko, Cene Fišer

**Affiliations:** Laboratory of Evolutionary Ecology of Hydrobionts, Nature Research Centre, Vilnius, Lithuania; SubBio Lab, Biotechnical Faculty, University of Ljubljana, Ljubljana, Slovenia

**Author notes:** Correspondence: Denis Copilaş-Ciocianu, Laboratory of Evolutionary Ecology of Hydrobionts, Nature Research Centre, Akademijos 2, 08412, Vilnius, Lithuania.

**Keywords:** Amphipoda, Cenozoic, climatic cooling, ecological radiation, molecular phylogeny, tectonic reconfiguration

## Abstract

The ecological radiation of amphipods is striking among crustaceans. Despite high diversity, global distribution and key roles in all aquatic environments, little is known about their ecological transitions, evolutionary timescale and phylogenetic relationships. It has been proposed that the amphipod ecological diversification began in the Late Palaeozoic. By contrast, due to their affinity for cold/oxygenated water and absence of pre-Cenozoic fossils, we hypothesized that the ecological divergence of amphipods arose throughout the cool Late Mesozoic/Cenozoic. We tested our hypothesis by inferring a large-scale, time-calibrated, multilocus phylogeny, and reconstructed evolutionary patterns for major ecological traits. Although our results reveal a Late Palaeozoic amphipod origin, diversification and ecological divergence ensued only in the Late Mesozoic, overcoming a protracted stasis in marine littoral habitats. Multiple independent post-Jurassic radiations took place in deep-sea, freshwater, terrestrial, pelagic and symbiotic environments, usually postdating deep-sea faunal extinctions, and corresponding with significant climatic cooling, tectonic reconfiguration, continental flooding, and increased oceanic oxygenation. We conclude that the profound Late Mesozoic global changes triggered a tipping point in amphipod evolution by unlocking ecological opportunities that promoted radiation into many new niches. Our study also provides a solid, time-calibrated, evolutionary framework to accelerate research on this overlooked, yet globally important taxon.

## Introduction

Global environmental changes shaped biodiversity patterns throughout Earth’s history (Roelants *et al.* 2007; Hannisdal & Peters 2011; Condamine *et al.* 2013). Understanding the historical factors that triggered large-scale evolutionary radiations or extinctions remains a central tenet in evolutionary biology. Investigating the effects of these past changes at the planetary level requires suitable model systems which can be represented by species rich taxonomic groups with a global distribution and high ecological diversity.

The Amphipoda is among the most ecologically diverse and speciose crustacean orders, encompassing over 10,000 species (Arfianti *et al.* 2018; Horton *et al.* 2019) inhabiting all aquatic environments worldwide, from hadal depths to alpine freshwater streams, from lightless groundwater to tropical forests, and from sea bottom sediments to the entrails of gelatinous plankton (Bousfield 1983; Barnard & Karaman 1991; Lowry & Myers 2017). Amphipods are highly abundant and have an important function in structuring aquatic communities (Oliver *et al.* 1982; Duffy & Hay 2000; González *et al.* 2008; Best & Stachowicz 2014). Furthermore, due to their omnivorous diet and intermediary trophic position, they represent a key link between trophic levels, thus playing an essential role in nutrient recycling (Dangles & Malmqvist 2004; Piscart *et al.* 2011; Machado *et al.* 2019). The dispersal abilities of amphipods are poor due to egg brooding, lack of free-swimming larvae and extended parental care (Barnard & Karaman 1991; Thiel 1999; Väinölä *et al.* 2008). Consequently, populations can easily become genetically isolated, leading to high species diversity and biogeographical patterns which accurately reflect ancient historical events (Finston *et al.* 2007; Hou *et al.* 2011; Bauzà-Ribot *et al.* 2012; Copilaş-Ciocianu & Petrusek 2017; Copilaş-Ciocianu *et al.* 2019). Lastly, amphipods are emerging model organisms for research on development, regeneration, ecotoxicology and evolutionary biology (Fišer 2012; Weston *et al.* 2013; Kao *et al.* 2016; Naumenko *et al.* 2017; Fišer *et al.* 2018).

Despite global distribution, high abundance, ecological significance, and importance as emerging model organisms, only little is known about the evolutionary history of amphipods and the factors that triggered their impressive radiation. This scarcity of knowledge is due to several critical factors. Deep evolutionary relationships within the Amphipoda are uncertain. The most comprehensive phylogenetic studies were based on morphology (Lowry & Myers 2013, 2017), which is known to be highly homoplastic in amphipods (Berge *et al.* 2000). Indeed, molecular phylogenies at lower taxonomic levels do not fully agree with the morphology-based systematics (Macdonald *et al.* 2005; Havermans *et al.* 2010; Hurt *et al.* 2013; Esmaeili-Rineh *et al.* 2015; Hou & Sket 2016; Mamos *et al.* 2016; Verheye *et al.* 2016; Copilaş-Ciocianu *et al.* 2019). The age of the order is puzzling as well. It has been suggested that amphipods already appeared in the Late Palaeozoic, when the lineages of the superorder Peracarida split (Bousfield 1977, 1978; Schram 1986). Yet, unlike the rest of peracaridan orders with pre-Cenozoic fossil record (Schram 1986; Wolfe *et al.* 2016), the handful of amphipod fossil taxa are dated no earlier than Eocene and usually bear a modern appearance (Derzhavin 1927; Bousfield & Poinar 1994; Coleman 2006; Kupryjanowicz & Jażdżewski 2010). Hence, the temporal origin and main cladogenetic events of modern amphipods are not known and cannot be paralleled to main global environmental changes.

The present view on the modern diversity of amphipods can be summarised by two competing hypotheses. The first hypothesis, stating that most of the modern diversity has been attained by the end of the Palaeozoic or Early Mesozoic, is based on the current distribution of superfamilies in relationship to geochronology, cladistic relationships, or on patterns observed in related malacostracans (Barnard & Barnard 1983; Lowry & Myers 2013). The alternative hypothesis suggests that the diversity of amphipods could be much younger due to morphological continuity among higher taxa, and due to lower taxonomic diversity in terrestrial and deep-sea habitats in comparison to the closely related isopods (Bousfield 1978). Molecular phylogenetic studies tend to support this view because they indicate that the onset of diversification of several major amphipod clades dates to the Cretaceous/Palaeogene (Hou *et al.* 2014; McInerney *et al.* 2014; Corrigan *et al.* 2014; Verheye *et al.* 2017; Copilaş-Ciocianu *et al.* 2019). Apart from the recent fossil record and molecular timetrees, several other independent lines of evidence point out to a more recent radiation of amphipods. Amphipods are particularly cold adapted animals, exhibiting an inverse latitudinal richness gradient in marine and freshwater habitats (Barnard 1976; Barnard & Barnard 1983; Barnard & Karaman 1991; Väinölä *et al.* 2008; Rivadeneira *et al.* 2011; Copilaş-Ciocianu *et al.* 2019), and high diversity and dominance in the cold, deep-sea benthic assemblages (De Broyer *et al.* 2004; Verheye *et al.* 2017; Havermans & Smetacek 2018). This pattern is probably related to their generally low tolerance to hypoxia, given that warmer water has a lower concentration of dissolved oxygen (Modig & Ólafsson 1998; Wiklund & Sundelin 2001; Wu & Or 2005; Vaquer-Sunyer & Duarte 2008). As such, it seems unlikely that amphipods could have attained most of their current ecological disparity during the Early to Middle Mesozoic (Triassic to Early Cretaceous), a period characterized by warm temperatures even in the deep-sea, by weakly stratified oceans and frequent anoxic events that caused major extinctions (Lear *et al.* 2000; Jacobs & Lindberg 2002; McClain & Hardy 2010). Therefore, we hypothesize that amphipods ecologically radiated in the Late Mesozoic/Cenozoic, when large-scale continental reconfiguration induced global climatic cooling, causing the oceans to transition to a thermohaline (two-layered) circulation which, in turn, increased productivity and oxygenation levels (McClain & Hardy 2010; Donnadieu *et al.* 2016; Mills *et al.* 2019). To test our hypothesis, we generated the first large-scale, time calibrated molecular phylogeny of the Amphipoda and reconstructed the course of diversification and ecological transitions. We focused on the timing of major cladogenetic events and ecological transitions.

## Materials and Methods

### Data collection and sequence alignment

As a taxonomic backbone for data collection we used the classification on the World Register of Marine Species database (WoRMS; http://www.marinespecies.org/) which is based on Lowry & Myers (2017). All the data used in the present study is publicly available in GenBank (www.ncbi.nlm.nih.gov/genbank) and originates from 63 published and 13 unpublished studies (a list of the data sources is found in Appendix 1 and Table S1; data collection ended in January 2018). Taxa were included in a way that we would cover as much phylogenetic and ecological diversity as possible. Maximizing phylogenetic diversity diminishes the effect of long-branch attraction and increases topological accuracy by dispersing homoplasy across the tree (Heath *et al.* 2008). In most cases, we included one representative species per genus. We selected four molecular markers based on their abundance and representativeness for all main clades: the mitochondrial cytochrome c oxidase subunit I (COI), the nuclear ribosomal RNA for the large and small subunits (28S and 18S), and the nuclear histone 3 (H3). All sequences were screened for contamination, presence of stop codons and homology. Preliminary gene trees were constructed for each marker to identify and eliminate unreliable sequences. To properly root the phylogeny, we included nine outgroups representatives: the sister order Ingolfiellida, as well as other members of the Peracarida and Decapoda. The dataset contained 210 (201 ingroup) terminals, representing 102 of the 226 recognized families (45%) (Table S1).

The PhyRe python script (Plazzi *et al.* 2010) was used to assess the phylogenetic representativeness of our dataset. The analysis was run at the genus level and the reference taxonomy was obtained from the WoRMS database. Confidence intervals for the average and the variation in taxonomic distinctness were calculated using 1000 random replicates.

The protein coding COI and H3 sequences were aligned with MUSCLE (Edgar 2004) in MEGA 6 (Tamura *et al.* 2013) and amino acid translated to check for premature stop codons (indicating pseudogenes). Following Copilaş-Ciocianu *et al.* (2018), we separately aligned the 18S and 28S rRNA sequences with SATé 2.2.7 (Liu *et al.* 2012). SATé simultaneously co-estimates the alignment and phylogenetic tree, which makes it far more accurate than other alignment methods (Mirarab *et al.* 2015). MAFFT 6.7 (Katoh *et al.* 2005) was used as the aligner and OPAL 1.0.3 (Wheeler & Kececioglu 2007) as the merger because this combination provides the highest phylogenetic accuracy (Liu & Warnow 2014). For tree inference we used the maximum-likelihood method implemented in FastTree 2.1.4 (Price *et al.* 2010) with the GTR+G20 substitution model. The cycle of alignment and tree building was repeated ten times for each marker. The alignments with the best likelihood score were used as input for final tree estimation and statistical support analyses (see *Phylogenetic reconstruction*). Gblocks 0.9 (Talavera & Castresana 2007) was used to remove poorly aligned regions with questionable homology in the 18S and 28S alignments. Minimum restrictive settings were applied and regions with gap positions were allowed in the final alignment. The final alignment length was 1741 bp for 18S, 883 bp for 28S, 436 bp for COI (third codon position removed; see next section), and 327 bp for H3, totalling 3387 bp. Individual marker alignments were concatenated using Sequence Matrix (Vaidya *et al.* 2011). The degree of missing data was 29%. The alignment in NEXUS format is available on Figshare (doi available after acceptance).

Ecological data regarding habitat (marine, freshwater and terrestrial; benthic vs. pelagic), mode of life (free vs. symbiotic), depth (littoral/epipelagic, shelf/mesopelagic, bathyal/bathypelagic, abyssal/abyssalpelagic and hadal/hadalpelagic) and temperature (cold vs. warm) were gathered from the relevant literature at the genus level (Bazikalova 1945; Laval 1980; Barnard & Barnard 1983; Barnard & Karaman 1991; Vinogradov *et al.* 1996; de Broyer *et al.* 2007). Note that “symbiotic mode of life” encompasses different types of symbiosis (commensalism, parasitism, and amensalism). The depth zone was attributed on the mean depth value, obtained from the minimum and maximum values for depth ranges of each taxon. A taxon was considered as cold water distributed when its representatives occurred at high to temperate latitudes or deeper than 1000 m. Similarly, a taxon was classified as warm water when its representatives were distributed in tropical to temperate waters above 1000 m depth. All ecological data can be found in the supplementary information (Table S2). To provide an overview of the geographic distribution of clades, we also gathered distribution data which was obtained from Barnard & Karaman (1991). Terminals were assigned to 17 geographical areas (Table S2).

### Phylogenetic reconstruction

We evaluated the level of substitution saturation of each marker using the index of Xia et al. (2003) implemented in DAMBE 5.3.10 (Xia & Xie 2003). Significant levels of saturation were detected at the COI 3^rd^ codon position (I_ss_>I_ss.c_, *p* = 0.0001), and, as such, these sites were not included in the phylogenetic analyses. Variable and parsimony informative sites were calculated in MEGA. The concatenated alignment contained 1917 parsimony informative out of 2333 variable sites (18S: 1006/1273; 28S: 575/667; COI: 209/251; H3: 127/142). Best-fitting evolutionary models were selected using PartitionFinder 2 (Lanfear *et al.* 2017) under the Bayesian Information Criterion and greedy search option.

Phylogenetic relationships were inferred using maximum likelihood (ML), Bayesian inference (BI) and maximum parsimony (MP) methods. The ML analyses were conducted with IQTREE 1.6 (Nguyen *et al.* 2015) and RAxML HPC 8.2.10 (Stamatakis 2014). The IQTREE search was performed under an edge-linked partitioned model (applied to each gene partition), using the GTR model with free rate heterogeneity (+R) which relaxes the assumption of Gamma distributed rates and has a better fit to large and complex datasets (Yang 1995; Soubrier *et al.* 2012). Statistical support for branches was assessed using 1000 ultra-fast bootstrap replicates (UFBS; Hoang et al. 2018) and the Shimodaira-Hasegawa approximate likelihood ratio test (SH-aLRT; Shimodaira and Hasegawa 1999, Guindon et al. 2010). The RAxML analysis was run with the GTR+Γ model applied to each gene partition. A thorough ML search was performed and 1000 rapid bootstrap replicates (RBS) were used to assess branch support. Bayesian analyses were performed with ExaBayes 1.5 (Aberer *et al.* 2014) under the GTR+Γ model applied to each gene partition. All parameters (except branch length) were unlinked and rates were allowed to vary independently. The analysis was run for 10^7^ iterations, with a thinning of 500 and 50% burn-in. The value for the parsimony subtree pruning and regrafting (SPR) radius parameter was set to 50 and the number of swaps per generation to 10. Maximum parsimony was performed with PAUP* 4.0a164 (Swofford 2002), using heuristic searches with TBR branch swapping and 1000 random taxon additions. Only phylogenetically informative sites were retained, gaps were treated as missing data, and all characters were unordered and equally weighted. Nodal support was estimated with 500 jackknifing replicates (JK) with 50% character removal. All phylogenetic analyses were performed on the CIPRES Science Gateway v3.3 (Miller *et al.* 2010).

### Molecular dating

Molecular dating was performed in BEAST 1.8.2 (Drummond *et al.* 2012) using the GTR+I+Γ model for COI and H3, and SYM+I+Γ for 18S and 28S (as selected with PartitionFinder). The ML phylogram from the IQTREE analysis was used as a starting tree in order to reduce computational time. An uncorrelated relaxed clock with a lognormal distribution was applied to each partition and speciation was modelled using the Birth-Death process. The MCMC chain was run for 10^8^ generations, with a sampling frequency of 2000. Convergence of parameters and effective sample size were assessed with Tracer 1.6 (Rambaut *et al.* 2014) after discarding 20% of trees as burn-in. We performed three independent runs, which gave the same result. As such, all runs were combined using LogCombiner 1.8.2 and the maximum clade credibility tree was produced with TreeAnnotator 1.8.2, both part of the BEAST package.

For divergence times estimation we employed the fossil calibration scheme described in detail by Copilaş-Ciocianu *et al.* (2019), to which we added one more calibration point. Only fossil taxa that are well studied and represented by several specimens were used for calibration. All calibration nodes were assigned exponential prior distributions since they require fewer parameters and are more appropriate when the fossil record of the focal group is poorly known (Ho & Phillips 2009). Briefly, we used five calibration points, the youngest (node 1), representing the origin of the Ponto-Caspian gammarid amphipod radiation, was set to a minimum age of 9 Ma (mean = 25, offset = 8, 95% HPD = 9–83) based on Caucasian fossil specimens (Derzhavin 1927). The Niphargidae/Pseudoniphargidae and Crangonyctidae/Pseudocrangonyctidae splits (nodes 2 and 3, respectively) were set at a minimum of 35 Ma (mean = 60, offset = 35, 95% HPD = 38–215) based on Eocene Baltic amber fossils (Coleman & Myers 2000; Coleman 2004, 2006; Kupryjanowicz & Jażdżewski 2010). The additional calibration point (node 4) that we use in this study is based on Miocene amber specimens of the family Talitridae from Central America (Bousfield & Poinar 1994, 1995). The minimum age was set to 25 Ma (mean = 30, offset = 21, 95% HPD = 22–111) and was applied to the stem of the Talitridae because this family has an uncertain phylogenetic position within the Talitroid clade of our phylogeny. Finally, the oldest calibration point (node 5) was placed as close to the root as possible (following Duchêne *et al.* 2014), and represents the oldest known member of Eumalacostraca, *Palaeopalaemon newberry* Whitfield, 1880 (minimum = 358 Ma, mean = 55, offset = 355, 95% HPD = 358–514) (Schram *et al.* 1978). The nodes do not seem misdated, given that inconsistency between fossil ages and lineage history was not significant (Shapiro-Wilk normality test, W = 0.91, *p* = 0.51, see Marshall 2008).

Due to the notable difference between the recent amphipod (ingroup) calibration points (9-35 Ma) and the old outgroup (358 Ma) calibration, we also ran two additional analyses, one only with ingroup calibrations, and one only with the outgroup calibration to assess if they produce compatible results.

### Diversification through time and ancestral state reconstruction

A sliding window analysis was performed according to Meredith *et al.* (2011) to visually inspect the tendency of diversification rates through time. The period between 180 and 10 Ma was divided into sliding windows of 10 Ma, with a frequency of 2.5 Ma. The period prior to 180 Ma was not considered due to the low number of lineages which would indicate an artificially inflated diversification rate. The number of lineages originating in a particular sliding window was divided by the number of lineages occurring prior to the start of that respective sliding window.

We reconstructed ancestral states for the five abovementioned ecological traits with the aim of evaluating the temporal framework of the amphipod ecological transitions. As such, we used the time-calibrated tree from the BEAST runs in subsequent analyses. Traits were treated as discrete and analysed with the re-rooting method using maximum likelihood (ML) (Yang *et al.* 1995) as well as stochastic character mapping (SCM) using Bayesian inference (Bollback 2006), both implemented in R (v.3.5.2) package *phytools* (v.0.6-60) (Revell 2012; R Core Team 2018). ML ancestral state reconstruction was performed with the *rerootingMethod* function, while SCM with the *make.simmap* function (Revell 2012). In order to estimate the possible ancestral character states on internal nodes, we ran 1000 simulations of stochastic character histories, using continuous-time reversible Markov model parameters of trait evolution, estimated using default settings and the character states on the tips of the phylogeny.

In order to evaluate the tempo of ecological disparification through time, we divided the phylogenetic history in 10 MY time bins, and calculated the average number of all possible transitions between states per time bin, as well as for the whole phylogeny. For the sake of clarity, we refer the term “diversification” to the process of speciation, while we use the term “disparification” to the process of ecological divergence.

## Results

### Dataset

The phylogenetic representative analysis indicated a highly representative taxon sampling. The Average Taxonomic Distinctiveness (AvTD) and Variation in Taxonomic Distinctiveness (VarTD) were above the highest AvTD and below the mean VarTD respectively (Fig. S1). Furthermore, von Euler’s index of imbalance (*I*_*E*_=0.102) was well below the recommended 0.25 threshold value, indicating unbiased sampling (Plazzi *et al.* 2010) (Fig. S1).

### Phylogenetic reconstruction and molecular dating

All five phylogenetic reconstruction methods yielded congruent results by recovering the same major clades. All model based methods recovered similar topologies and discordance was observed mainly at poorly supported nodes (Fig. 1, Figs. S2-S5). Altogether, we identified nine major clades which we named informally: Gammaroids– mainly northern hemisphere taxa, with freshwater proclivity, Lysianassoids–mostly deep-sea scavengers, Crangonyctoids– Holarctic freshwater species, Corophioids– tube-building epifaunal/infaunal marine species, Eusiorids/Iphimedioids– ecologically diverse cold-water marine species, Physosomatans and Physocephalatans–exclusively commensal and parasitic warm-water marine species, most of them pelagic, Talitroids–shallow water, ecologically diverse containing the only terrestrial lineage, and Atylids–a morphologically plesiomorphic and cosmopolitan marine group. The following groups were either weakly supported or polyphyletic, but we retained them for the sake of brevity: the Miscellaneous clade was weakly supported and comprised morphologically and ecologically disparate families, while the Hadzioids were morphologically consistent but polyphyletic. The order Ingolfiellida was recovered as a sister to Amphipoda only in the IQTREE analysis. All remaining analyses (ML with RAxML, BI, and MP) recovered the order Spelaeogriphacea as a sister to amphipods with high support, while the order Ingolfiellida was sister to (Amphipoda+Speleogriphacea) clade (Figs. S3-S5).

**Fig. 1.**
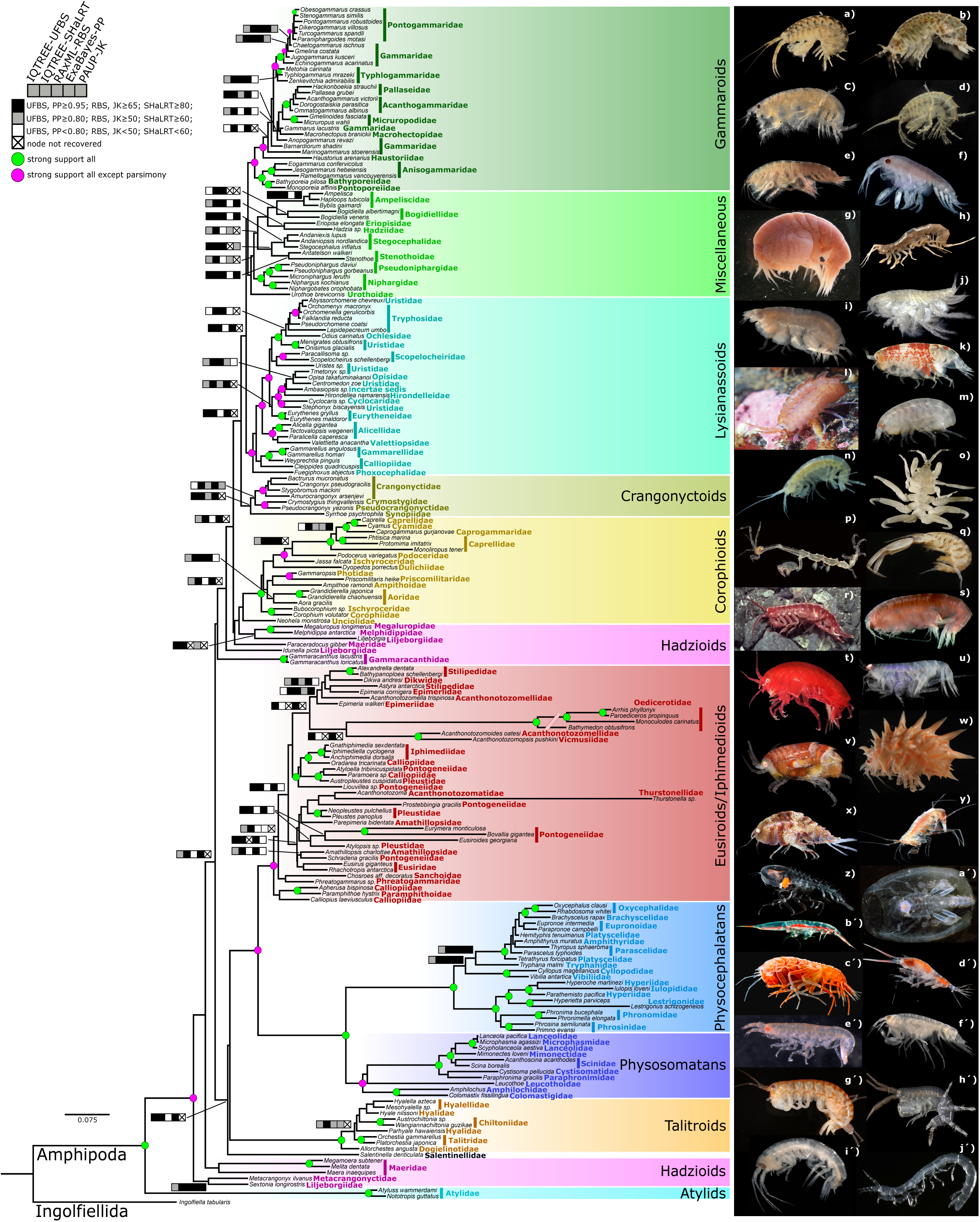
Maximum likelihood phylogram obtained from the concatenated dataset (COI, H3, 28S and 18S) using IQTREE. Nodes receiving strong support in all analyses are indicated with green circles, while nodes that received strong support in all analyses except parsimony are indicated with purple circles. Bars indicate the strength of the support that each node received in each analysis. Nodes that were weakly supported in all analyses are not annotated. Families are indicated with coloured font. Photographs on the right indicate the extent of morphological diversity of the Amphipoda. Photographs: Denis Copilas Ciocianu: a) *Dikerogammarus villosus*, b) *Chaetogammarus warpachowskyi*, d) *Gammarus roeselii*, h) *Niphargus pannonicus*; Dante Fenolio (used with author’s permission, photos available at www.anotheca.com/photograph): n) *Stygobromus* sp., z) *Cystisoma* sp., a’) *Phronima* sp., b’) *Streetsia* sp., c’) *Scypholanceola* sp., d’) *Scina* sp.; David Fenwick (used with author’s permission, photos available at www.aphotomarine.com): m) *Lysianassa ceratina*, p) *Caprella tuberculata*, q) *Corophium volutator*, u) *Pontocrates arenarius*, g’) *Orchestia gammarellus*, h’) *Melita hergensis*, j’) *Ingolfiella britannica*; Hans Hillewaert (licensed under Creative Commons, photos available at www.flickr.com/photos/bathyporeia/albums/72157639365477036): c) *Bathyporeia pilosa*, e) *Ampelisca brevicornis*, f) *Stenothoe marina*, i) *Lepidepecreum longicorne*, j) *Urothoe brevicornis*, o) *Cyamus boopis*, s) *Megaluropus agilis*, f’) *Leucothoe incise*, i’) *Nototropis swammerdamei*, v) *Iphimedia nexa*; Russ Hopcroft (licensed under Creative Commons, photo available at https://en.wikipedia.org): y) *Eusirus holmi*; Joanna Legeżyńska (licensed under Creative Commons, photo available at www.marinespecies.org): g) *Stegocephalus inflatus*; Gustav Paulay (licensed under Creative Commons, photo available at https://calphotos.berkeley.edu/): e’) *Colomastix* sp.; Martin Rauschert (licensed under Creative Commons, photo available at www.marinespecies.org): r) *Paraceradocus* sp.; Alexander Semenov (used with author’s permission, photos available at www.coldwater.science): k) *Aristias timidus*, l) *Gammarellus homari*, t) *Acanthonotozoma* sp., x) *Pleustes panopla*; Cédric d’Udekem d’Acoz (licensed under Creative Commons, photo available at www.marinespecies.org): w) *Epimeria oxicarinata*;

Molecular dating using all the calibration points or only the root calibration resulted in similar estimations (10 to 20 Ma differences; Table S3). The calibration scheme that included only the recent ingroup fossils resulted in expectedly younger estimates, however, the 95% HPD intervals of all calibration schemes overlapped to some extent (Table S3). Altogether, these results contradict the previous views that the modern diversity of the Amphipoda was already established by the Late Palaeozoic/Early Mesozoic (Barnard & Barnard 1983; Lowry & Myers 2013). For evolutionary, ecological and biogeographical interpretation we considered the complete calibration scheme since it is the most balanced and informed. Accordingly, Amphipoda has split off from Ingolfiellida during the Permian (∼281 Ma) and started radiating shortly after the Permo-Triassic mass extinction (∼240 Ma). Crown ages of all major clades lie between the Early Cretaceous and Early Palaeogene (Fig. 2, Table S3). A fully annotated chronogram is available as a supplementary figure (Fig. S6). An additional tree with the geographical distribution of taxa mapped onto it is also available as supplementary information (Fig. S7).

**Fig. 2.**
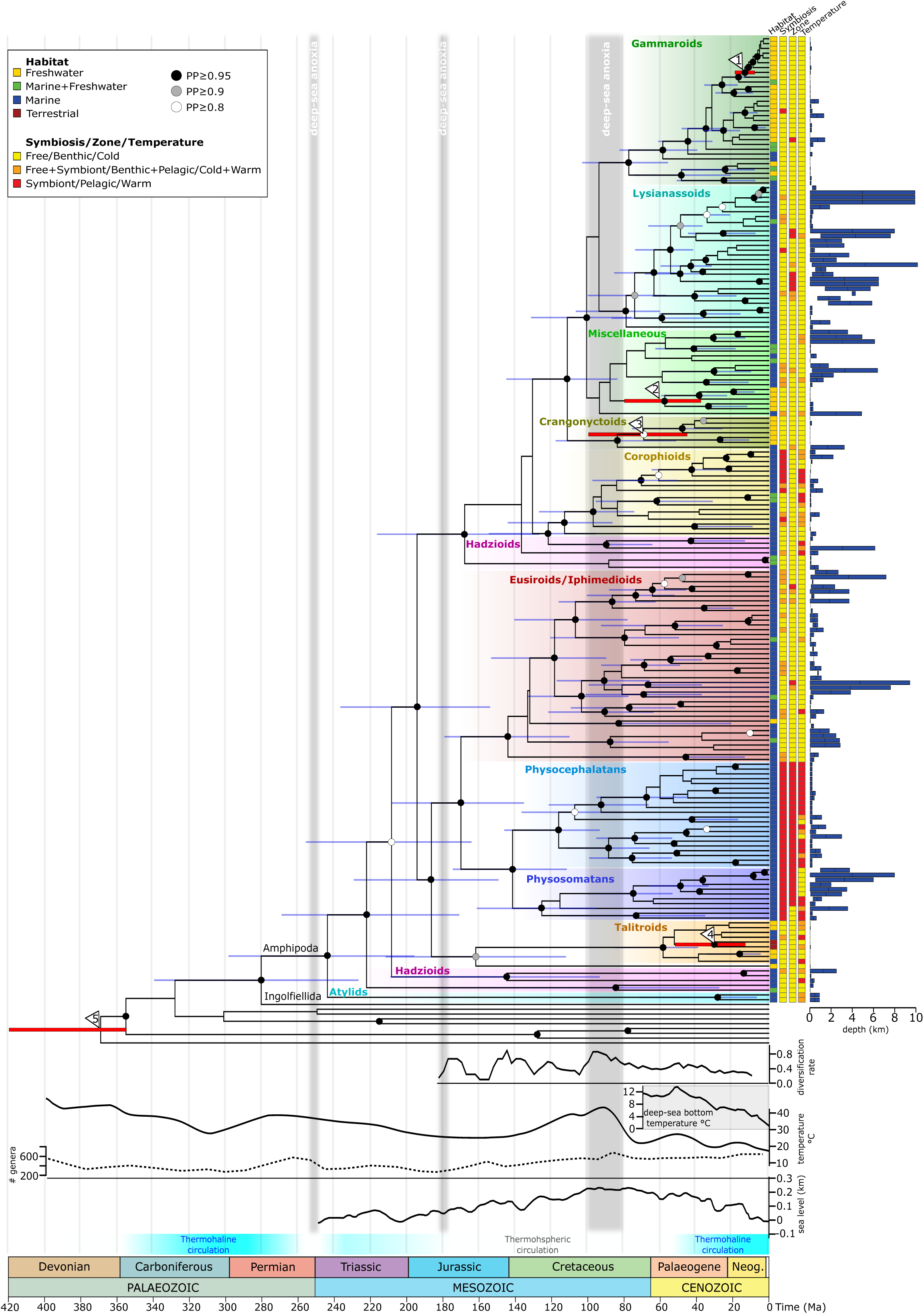
Evolutionary timescale of the amphipod radiation. The time-calibrated phylogeny was obtained with BEAST using the concatenated dataset (COI, H3, 28S and 18S). Circles at nodes indicate posterior probabilities (black ≥ 0.95, grey ≥ 0.9, white ≥ 0.8, not shown if < 0.8) and blue bars indicate the 95% HPD intervals of clade age (shown only for main and/or strongly supported nodes). Red bars accompanied by numbers indicate node ages that were constrained using fossils. The fully annotated chronogram can be found in the supplementary information (Fig. S6). Coloured boxes on the right summarize the ecological traits of each terminal in the tree (see legend in the top left) and bars indicate depth ranges. The line graphs below the phylogeny indicate (from top to bottom) the amphipod diversification rate through time, the evolution of Cenozoic deep-sea bottom temperature (Lear *et al.* 2000), the evolution of Phanerozoic surface temperature (Mills *et al.* 2019), the evolution of Phanerozoic invertebrate diversity (Alroy *et al.* 2008), and the evolution of the global Mesozoic-Cenozoic sea level (Miller *et al.* 2005). The gradient between turquoise and white colours indicates the transitions between thermohaline (two-layered ocean) and thermospheric (weakly stratified) circulation (McClain & Hardy 2010).

### Diversification through time and ancestral states reconstruction

The sliding window analysis identified four main pulses of diversification: the first occurred during the middle Jurassic, the second during the late Jurassic/early Cretaceous, the third during the mid-Cretaceous, and the fourth in the late Cretaceous (Fig. 2).

The SCM (Fig. 3) and ML (Fig. S8) reconstructions of ancestral states were highly congruent and revealed a dynamic evolution of Amphipod ecology. The analyses based on 1000 simulations suggested that all ecological changes, except the shift to semiterrestriality, happened multiple times: on average 5.7 times from free-living to symbiotic life-style, 7.4 times from marine to freshwater and 2.6 times vice versa, 12.9 times from cold to warm waters and 5.3 times back to cold water, and 9.5 times from benthic to pelagic habitat (Table 1). At least 100 changes of depth zones were estimated, mostly from shallow waters to deep sea (Table 1). There was a presumably single shift to semiterrestrial life from ancestors with uncertain salinity preference (Table 1, Fig. 3) (see Discussion).

**Table 1:** Average number of transitions (first column) and average number of transitions per time bin, from 1000 trees with mapped discrete character states. Cells with average changes ≥ 0.5 per time bin or average total change ≥1 are highlighted and maximum values per transition are bolded.

|  |  | Trias. | Jurassic |  | Cretaceous |  |  | Paleogene |  | Neo. |
| --- | --- | --- | --- | --- | --- | --- | --- | --- | --- | --- |
| Time bins | Mean | >200 | 200-170 | 170-140 | 140-110 | 110-80 | 80-60 | 60-40 | 40-20 | 20-0 |
| MODE OF LIFE |  |  |  |  |  |  |  |  |  |  |
| free - symbiont | 5.9 | 0 | 0 | 1 | 0.1 | 0.9 | 0.3 | 0.5 | 0.6 | 2.6 |
| symbiont - free | 0.2 | 0 | 0 | 0 | 0 | 0 | 0 | 0 | 0 | 0 |
| HABITAT |  |  |  |  |  |  |  |  |  |  |
| freshwater - marine | 2.6 | 0 | 0 | 0 | 0 | 0 | 0 | 0.2 | 1.5 | 0.8 |
| freshwater - semiterrestrial | 1 | 0 | 0 | 0 | 0 | 0 | 0 | 0.5 | 0.5 | 0 |
| marine - freshwater | 7.4 | 0 | 0.4 | 0.3 | 0.2 | 0.9 | 1.8 | 1.3 | 1.8 | 0.6 |
| marine - semiterrestrial | 0 | 0 | 0 | 0 | 0 | 0 | 0 | 0 | 0 | 0 |
| semiterrestrial - freshwater | 0 | 0 | 0 | 0 | 0 | 0 | 0 | 0 | 0 | 0 |
| semiterrestrial - marine | 0 | 0 | 0 | 0 | 0 | 0 | 0 | 0 | 0 | 0 |
| benthic - pelagic | 9.5 | 0 | 0 | 0.2 | 1 | 0.7 | 0.5 | 1 | 3.1 | 3.1 |
| pelagic - benthic | 0.7 | 0 | 0 | 0 | 0.1 | 0.2 | 0.1 | 0.1 | 0.1 | 0.2 |
| TEMPERATURE |  |  |  |  |  |  |  |  |  |  |
| cold - warm | 12.9 | 0.1 | 0.1 | 1 | 0.7 | 0.8 | 1.3 | 1.6 | 3.1 | 4.2 |
| warm - cold | 5.3 | 0.1 | 0 | 0.1 | 0.1 | 0.4 | 0.9 | 1 | 1.2 | 1.5 |
| DEPTH |  |  |  |  |  |  |  |  |  |  |
| [<0.2 km] - [0.2-1 km] | 35.5 | 0.3 | 0.4 | 0.9 | 1.9 | 3.4 | 4.5 | 5.6 | 7.4 | 11 |
| [<0.2 km] - [1-4 km] | 5 | 0 | 0 | 0.1 | 0.2 | 0.4 | 1.1 | 1 | 0.9 | 1.3 |
| [<0.2 km] - [>4 km] | 0 | 0 | 0 | 0 | 0 | 0 | 0 | 0 | 0 | 0 |
| [0.2-1 km] - [1-4 km] | 19.6 | 0.1 | 0.1 | 0.1 | 0.5 | 1.4 | 2.7 | 3.7 | 4.5 | 6.5 |
| [0.2-1 km] - [>4 km] | 1.5 | 0 | 0 | 0 | 0 | 0 | 0.2 | 0.2 | 0.2 | 0.9 |
| [1-4 km] - [>4 km] | 5.5 | 0 | 0 | 0 | 0 | 0 | 0.1 | 0.4 | 1.9 | 2.9 |
| [0.2-1 km] - [<0.2 km] | 19.1 | 0.2 | 0.2 | 0.4 | 0.8 | 1.6 | 2.4 | 3.4 | 4.4 | 5.8 |
| [1-4 km] - [<0.2 km] | 1.7 | 0 | 0 | 0 | 0 | 0.1 | 0.1 | 0.3 | 0.5 | 0.7 |
| [1-4 km] - [0.2-1 km] | 12.5 | 0 | 0 | 0.1 | 0.2 | 0.6 | 0.8 | 2 | 3.2 | 5.6 |
| [>4 km] - [0.2-1 km] | 1.2 | 0 | 0 | 0 | 0 | 0 | 0 | 0 | 0 | 1.1 |
| [>4 km] - [1-4 km] | 1 | 0 | 0 | 0 | 0 | 0 | 0 | 0.1 | 0.3 | 0.5 |
| [>4 km] - [<0.2 km] | 0 | 0 | 0 | 0 | 0 | 0 | 0 | 0 | 0 | 0 |

**Fig. 3.**
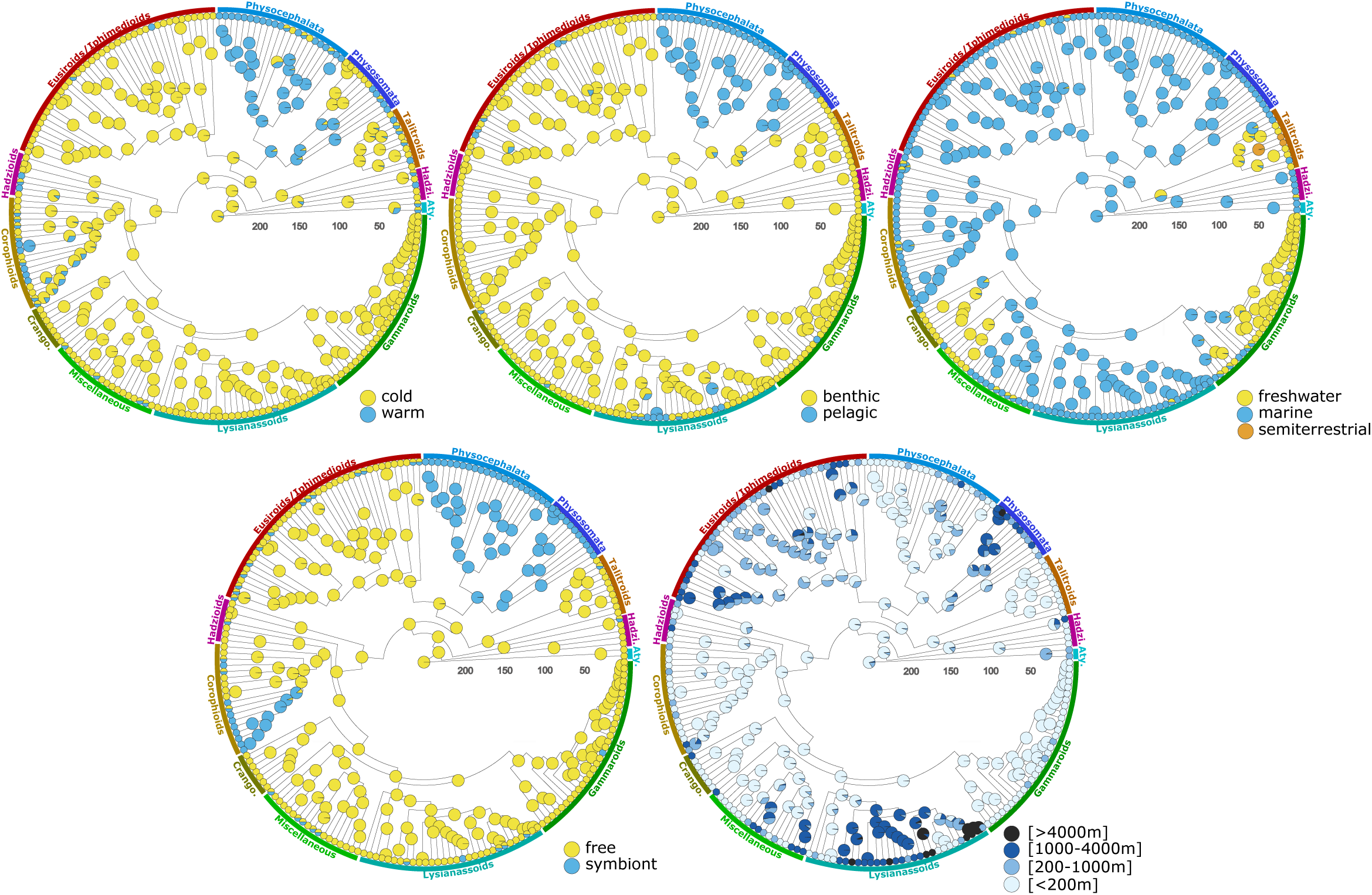
Ecological transitions of Amphipoda through time. Ancestral state reconstruction was estimated using stochastic character mapping. Pie-charts at nodes indicate the probability of ancestral states. The maximum likelihood results of the ancestral states are presented in the supplementary information (Fig. S8).

All these ecological changes took place relatively late in the amphipod evolutionary history, no earlier than 170Ma. The periods of intense ecological disparification corresponded to peaks of lineage diversification. The ancestral amphipod was reconstructed as a cold-water, free-living, marine animal, inhabiting shallow benthos. The first ecological shifts date to middle Jurassic, when the ancestor of Hyperiid clade (Physosomatans+Physocephalatans) likely switched from cold-water and free-living benthic species to warm-water and mostly pelagic symbiont mode of life (Table 1, Figs. 3-S8). The ancestral salinity of Talitroids is uncertain because the ML analysis suggests a marine ancestry while the SCM indicates a freshwater ancestry (see Discussion for details). The next pulse of disparification took place in Late Cretaceous, when the caprellid lineage of the Corophioid clade switched from cold-water free-living animals to warm water-symbionts. At the same time, at least two transitions to freshwater occurred in Crangonyctoids and Niphargidae/Pseudoniphargidae clades. Gammaroids presumably transitioned to fresh waters at least twice during the Palaeogene. Similarly, transition to deep-sea niches (>1000 m) occurred predominantly during the Palaeogene, mainly in the Lysianassoids, the Physosomatans and several lineages of the Eusiroid/Iphimediod clade. The third pulse of ecological disparification took place during the last 50 Ma, with multiple ecological shifts all over the phylogeny (Table 1, Fig3. 3-S8).

## Discussion

Our study reveals an ancient Permian origin of amphipods, and their delayed diversification during the Late Jurassic-Early Cretaceous followed by ecological disparification during the Cretaceous-Palaeogene. These results refute the view that most of the modern amphipod diversity already existed since the Late Palaeozoic-Early Mesozoic and help reconcile their old history with their absence in the pre-Cenozoic fossil record. Below we discuss the main global events that may have affected their diversification and ecological radiation. We focus on large-scale patterns and emphasize that although many nested radiations likely occurred discussing them in detail is beyond the scope of this study.

### Global evolutionary and biogeographical patterns

Our results indicate that amphipods split off from ingolfiellids during the Permian (∼280 Ma). This is in good agreement with previous studies which suggested a Late Palaeozoic age of the Amphipoda based on the fact that peracarids went through an extensive phase of radiation and dispersal during this time (Bousfield 1983; Schram 1986; Wolfe *et al.* 2016). Furthermore, the Permian is also known for its peak in the diversification of marine invertebrates (Alroy *et al.* 2008). Extant amphipod lineages started radiating after the major Permo-Triassic extinction, a catastrophic event that wiped out up to 95% of marine taxa (Benton & Twitchett 2003). During this dramatic extinction, the Panthalassic Ocean was largely anoxic, especially at higher latitudes or depths, and only the Palaeotethys Ocean retained suitable levels of oxygenation (Penn *et al.* 2018), where hypoxia-sensitive amphipods could survive and from where they subsequently radiated. This hypothesis is concordant with tethyan distribution of the basal, shallow-water Hadziod lineages in our phylogeny (Barnard 1976; Stock 1993; Bauzà-Ribot *et al.* 2012).

Most of the deep splits in the phylogeny leading towards the extant major clades took place during the Triassic and Early Cretaceous (Fig. 2). Also, three of the four main lineage diversification peaks took place in this period. Assuming that amphipods were littoral inhabitants throughout this time-frame, it is likely that the breakup of Pangaea followed by subsequent fragmentation of the resulting Laurasia and Gondwana supercontinents (Seton *et al.* 2012) played an important vicariant role, as seen in the Holarctic freshwater crangonyctoids (Copilaş-Ciocianu *et al.* 2019) or thalassoid lineages with an amphi-Atlantic distribution (Bauzà-Ribot *et al.* 2012). We consider that more detailed inferences are speculative due to the cosmopolitan distribution of many marine lineages and the lack of pre-Cenozoic fossils. Exploring this issue further requires more data and in-depth phylogenetic and biogeographical analysis of the cosmopolitan clades. Similarly, a detailed analysis of patterns of diversification rates through time was not realistic due to the extremely scarce amphipod fossil record and our incomplete dataset lacking half of the known families (Marshall 2017; Diaz *et al.* 2019).

The last major spike in the amphipod diversification history occurred ca. 90 Ma ago (Fig. 2). This peak of diversification could be related to the globally high sea-level stands which resulted in vast shallow epicontinental seas that provided a lot of habitat suitable for amphipods (Scotese 2014). This is particularly visible in the warm, shallow-water Corophioid, Physocephalatan and Physosomatan clades which started proliferating in this period. Furthermore, this diversification peak coincides with a peak in diversity of marine invertebrates (Alroy *et al.* 2008) and also to the highest levels of atmospheric oxygen during the last 250 Ma (Berner 2002).

Our estimated time-frame of overall amphipod diversification agrees well with recent molecular studies which consistently indicate a Cretaceous-Palaeogene diversification of several major clades, despite different approaches to calibrate divergence times (Hou *et al.* 2014; McInerney *et al.* 2014; Corrigan *et al.* 2014; Verheye *et al.* 2017; Copilaş-Ciocianu *et al.* 2019). Therefore, these studies provide strong evidence to indicate that the diversity of extant amphipods is indeed relatively recent, despite the old Permian origin of the group.

### Ecological radiation

Our ancestral state reconstructions suggested that ancestral amphipods were free-living, marine, littoral animals with an affinity for cold water for a large part of their evolutionary history (Permian to Late Jurassic), and are in agreement with previous hypotheses (Bousfield 1983). The littoral, which is the most oxygenated part of the ocean (Keeling *et al.* 2010) possibly acted as a long-term refugium for hypoxia sensitive animals such as amphipods (Modig & Ólafsson 1998; Wiklund & Sundelin 2001; Wu & Or 2005; Vaquer-Sunyer & Duarte 2008). This would suggest that the long Permian-to-mid Mesozoic ecological stasis in littoral habitats was a result of environmental constraints, reflecting the suboptimal conditions present in non-littoral parts of the ocean, such as high global temperatures (especially in the deep sea), weak oceanic circulation, hypoxic conditions and frequent deep-sea anoxia (McClain & Hardy 2010; Mills *et al.* 2019). Considering that the coastal environment is prone to erosion rather than deposition (Wilke *et al.* 2016), our results support the view that the absence of a pre-Cenozoic amphipod fossil record (Bousfield 1978; Schram 1986) is best explained by a combination of long evolutionary confinement to active littoral habitats, small size and weakly fossilizable cuticle (Bousfield 1983).

The overall evidence thus indicates that the ecological radiation of amphipods proceeded from shallow marine waters. Most of the freshwater radiations emerged relatively recently (Late Cretaceous/Cenozoic) after the substantial Late Cretaceous temperature drop (Figs. 2-3)(Mills *et al.* 2019). All of these freshwater clades (especially the Holarctic Crangonyctoids, Gammaroids and Niphargids) descended from shallow-water ancestors and often exhibit a peculiar biogeographic pattern with older lineages and higher phylogenetic diversity occurring in the northern part of their ranges (McInerney *et al.* 2014; Copilaş-Ciocianu *et al.* 2017, 2019). This is likely a relict pattern which reflects their more northwards pre-Pleistocene distribution, and consequently their affinity for cold water. This hypothesis is supported by amber fossils clearly belonging to the Niphargidae clade, found in the Baltic region, which lies northwards of the contemporary range of the group (Coleman & Myers 2000; Kupryjanowicz & Jażdżewski 2010). Noteworthy, the ancestral salinity preference of the Talitroid clade is uncertain in our analyses. The Bayesian reconstruction indicated a freshwater ancestry (Fig. 3) while the ML reconstruction suggested a marine ancestry (Fig. S8). These contrasting results are most likely an artefact of low sampling of marine Talitroid lineages due to limited sequence availability. Most likely the group has a marine origin (Serejo 2004) and more fine scale studies also showed a saline origin of some inland Talitrids (Yang *et al.* 2013).

All the colonizations of the deep-sea also took place throughout the cooling of the Late-Cretaceous/Cenozoic, and postdate the Cenomanian-Turonian oceanic anoxic event which triggered global faunal extinctions (Figs. 2-3)(80-100 Ma; Arthur *et al.* 1987). This time frame also corresponds with the opening and deepening of the North Atlantic and Southern Oceans (Seton *et al.* 2012) which are thought to have played an important role in the diversification of the Lysianassoid and Eusiroid/Iphimedioid clades (Corrigan *et al.* 2014; Verheye *et al.* 2017). These relatively recent deep-sea radiations agree well with the prevailing view that most of the deep-sea fauna is of Cenozoic origin due to the hypoxic conditions during most of the Mesozoic (Lindner *et al.* 2008; McClain & Hardy 2010; Vrijenhoek 2013; Herrera *et al.* 2015). At least for deep-sea amphipods, the tectonic-induced climatic cooling, leading to a better ventilated ocean (Donnadieu *et al.* 2016), coupled with the Cenomanian-Turonian extinction event not only dwindled the competitors but also created suitable environmental conditions for amphipods to thrive and ecologically expand. Moreover, the inverse latitudinal richness gradient observed in contemporary freshwater and marine amphipods (Barnard & Barnard 1983; Väinölä *et al.* 2008; Rivadeneira *et al.* 2011) further emphasizes that low temperature is somehow critical for the colonization of freshwater and deep-sea niches. However, we lack a precise understanding of these processes, which are probably related to the generally low tolerance of amphipods to hypoxic conditions (Modig & Ólafsson 1998; Wiklund & Sundelin 2001; Wu & Or 2005; Vaquer-Sunyer & Duarte 2008).

Transitions to a pelagic life-style occurred numerous times as well, and are especially prevalent in deep-sea lineages (Fig. 3). In freshwater, this transition occurred only once, in the highly specialized Baikal species *Macrohectopus branickii*. Like in the previous cases, the majority of these pelagic transitions are relatively recent (Late Cretaceous to Cenozoic), mainly because these lineages occur in the deep-sea. Exceptionally, the diverse Physocephalatan clade colonized warm shallow epipelagic waters during the Early Cretaceous, earlier than all of the other pelagic lineages. Our results reveal that Hyperiids (Physocephalatans+Physosomatans) colonized pelagic niches two times independently and contradict the prevailing view of a single colonization of pelagial (Lowry & Myers 2017).

The expansion into symbiotic niches seems to not have been influenced that much by climatic cooling. Both the large symbiotic Hyperiid clade as well as the caprellid linage of the Corophioid clade switched to this life-style during the warm Late Jurassic/Cretaceous (Figs. 2-3). These clades are associated with an ancestral preference for warmer temperatures (Fig. 3). Therefore, it is likely that the Cretaceous high sea-level which led to the formation of numerous shallow and warm epicontinental seas (Scotese 2014) created vast amounts of suitable habitats which promoted diversification and facilitated the formation of symbiotic relationships. The crown age of the exclusively symbiotic Hyperiid clade (ca. 140 Ma) corresponds well with a peak of reef forming coral diversity at 150 Ma (Simpson *et al.* 2011), an important habitat for the basal members of this clade (Barnard & Karaman 1991; Lowry & Myers 2009). These symbiotic clades often contain taxa with extreme ecological specialization, and exhibit a highly modified morphology (Fig. 1) (Laval 1980; Ito *et al.* 2011; Hurt *et al.* 2013).

The colonization of terrestrial habitats occurred only in the lineage of the family Talitridae of the Talitroid clade. Although this is possibly the most radical ecological transition among amphipods (Spicer *et al.* 1987), it is also one of the most recent, occurring during the Palaeogene (Figs. 2-3). It has been previously proposed that the terrestrial transition of talitrids was dependent on the availability of angiosperms (main food source), and thus, should not be older than the Cretaceous radiation of this group of plants (Bousfield 1983; Barba-Montoya *et al.* 2018). Nevertheless, some authors suggest a Pangaean origin of talitrids given their global distribution (Lowry & Myers 2019). Additional evidence supporting a young age for this clade is its less advanced stage of terrestrial specialization and far lower taxonomic diversity in comparison with the related oniscoidean isopods (Bousfield 1983; Spicer *et al.* 1987), which have colonized this habitat at least since the Early Cretaceous, possibly even the Late Palaeozoic (Broly *et al.* 2013). The young age of the Talitridae implies that the switch to semiterrestiral life likely took place multiple times within the Talitroid lineage, a hypothesis that remains to be tested using more comprehensive taxon sampling.

### Systematic implications

The main molecular clades recovered in our phylogeny correspond well in most cases with the major morphological groups (Lowry & Myers 2017). Due to some unresolved basal nodes and incomplete taxon sampling, the topology of our molecular phylogeny cannot refute the current systematic view. Although we had data on four of the six recognized suborders (Hyperiopsidea and Pseudoingolfiellidea were missing), our phylogeny generally supports the three main suborders, Amphilochidea, Hyperiidea and Senticaudata, but suggests some reshuffling. For example, the parvorders Lysianassidira, as well as the polyphyletic Haustoriidira and Synopiidira should be placed into the Senticaudata rather than Amphilochidea. Similarly, the suborder Colomastigidea along with the amphilochidean families Amphilochidae and Leucothoidae should be placed within Hyperiidea, reinforcing a previously proposed close relationship between Hyperiidea and Amphilochidae (Kim & Kim 1993). We consider that a thorough systematic discussion is beyond the scope of this paper, and suggest that a phylogenomic approach could prove invaluable for a better understanding of amphipod evolution, as evidenced in a recent study on decapod crustaceans (Wolfe *et al.* 2019). However, any evolutionary hypothesis needs to be tested by independent lines of evidence, and, as such, we consider that phenotypic data plays a crucial role even in the modern era of phylogenomics (Lee & Palci 2015).

## Conclusion

The Late Mesozoic is notable for its dramatic global changes which saw the rise and demise of many organismal groups, leading towards the modern biota (Scott 1995; Roelants *et al.* 2007; Alroy *et al.* 2008; Schulte *et al.* 2010; Meredith *et al.* 2011; Barba-Montoya *et al.* 2018; Varga *et al.* 2019). In the case of the Amphipoda, these changes brought an important turning point in their evolution. The fortuitous coupling of several critical circumstances such as extinction of deep-sea competitors followed by climatic cooling, oceanic deepening and increased oxygenation, created ecological opportunities which allowed hypoxia-sensitive amphipods to overcome a long period of ecological stasis and to radiate into many new niches. Such patterns of protracted stasis followed by extensive ecological radiation due to global changes seem to be common throughout the tree of life (Mitchell & Makovicky 2014; Halliday *et al.* 2019). Our results also emphasize the importance of molecular phylogenetics in illuminating the evolutionary history of highly diverse clades which have an extremely poor fossil record.

## Acknowledgements

We thank Dante Fenolio, David Fenwick and Alexander Semenov for kindly providing amphipod photographs. DCC was supported by the Research Council of Lithuania (09.3.3-LMT-K-712-13-0150), šB and CF were supported by the Slovenian Research Agency (Program P1-0184, Project N1-0069 and grant contract. KB139 382597).

## Data accessibility

Trees and alignments will be available at FigShare and Treebase.

## Author contributions

DCC, CF and SB conceived the study, DCC collected data, performed phylogenetic and molecular dating analyses, SB performed ancestral state reconstructions and diversification analyses, DCC lead the writing, all authors contributed to and approved the final version of the manuscript.

## Appendix

Reference list of all studies which generated the sequence data used in the current work

## Supporting Information

Table S1 Sequence data used in this study and its origin

Table S2 Ecological and geographical data

Table S3 Comparison of clade ages under different calibration schemes

Fig. S1 Funnel plots resulting from the PhyRe analyses

Fig. S2 Maximum likelihood phylogeny obtained with IQTREE.

Fig. S3 Maximum likelihood phylogeny obtained with RAxML.

Fig. S4 Bayesian phylogeny obtained with ExaBayes

Fig. S5 Maximum parsimony phylogeny obtained with PAUP Fig. S6 Fully annotated Bayesian chronogram.

Fig. S7 Geographical distribution of taxa used in the phylogenetic analysis.

Fig. S8 Maximum-likelihood reconstruction of ancestral states.

