## Supplemental Data 1 for "The late blooming amphipods: global change promoted post-Jurassic ecological radiation despite Palaeozoic origin"

### Table of Contents:


| | |
|---|---|
| **Table S3** | Page 2 |
| **Fig. S1 Funnel plots resulting from the PhyRe analyses** | Page 3 |
| **Fig. S2 Maximum likelihood IQTREE phylogeny** | Page 4 |
| **Fig. S2 Maximum likelihood RAxML phylogeny** | Page 5 |
| **Fig. S4 Bayesian ExaBayes phylogeny** | Page 6 |
| **Fig. S5 Maximum parsimony PAUP phylogeny** | Page 7 |
| **Fig. S6 Fully annotated chronogram** | Page 8 |
| **Fig. S7 Geographical distribution of taxa used in the phylogenetic analysis** | Page 9 |
| **Fig. S8 Maximum-likelihood reconstruction of ancestral states** | Page 10 |

Table S3 Comparison of divergence times (median and 95% HPD intervals of crown ages) for major focal clades under different calibration schemes (parentheses indicate the number of calibration points).

| Major clade | All fossils (5) | Root only (1) | Ingroup only (4) |
|---|---|---|---|
| Gammaroids | 78 [55–103] | 85 [61–117] | 37 [23–57] |
| Lyssianasoids | 79 [59–104] | 90 [66–119] | 45 [35–63] |
| Crangonyctoids | 68 [43–95] | 80 [50–114] | 49 [34–72] |
| Corophioids | 98 [72–126] | 112 [83–148] | 62 [43–89] |
| Eusiroids/Iphimedioids | 141 [107–177] | 162 [125–207] | 86 [62–120] |
| Physocephalatans | 113 [87–141] | 127 [98–162] | 73 [51–101] |
| Physosomatans | 122 [89–156] | 134 [99–175] | 77 [55–110] |
| Talitroids | 59 [38–87] | 72 [47–105] | 40 [25–58] |
| Amphipoda | 240 [189–292] | 257 [204–320] | 142 [100–201] |
| Amphipoda+Ingolfiellida | 281 [225–338] | 298 [236–373] | 174 [117–252] |

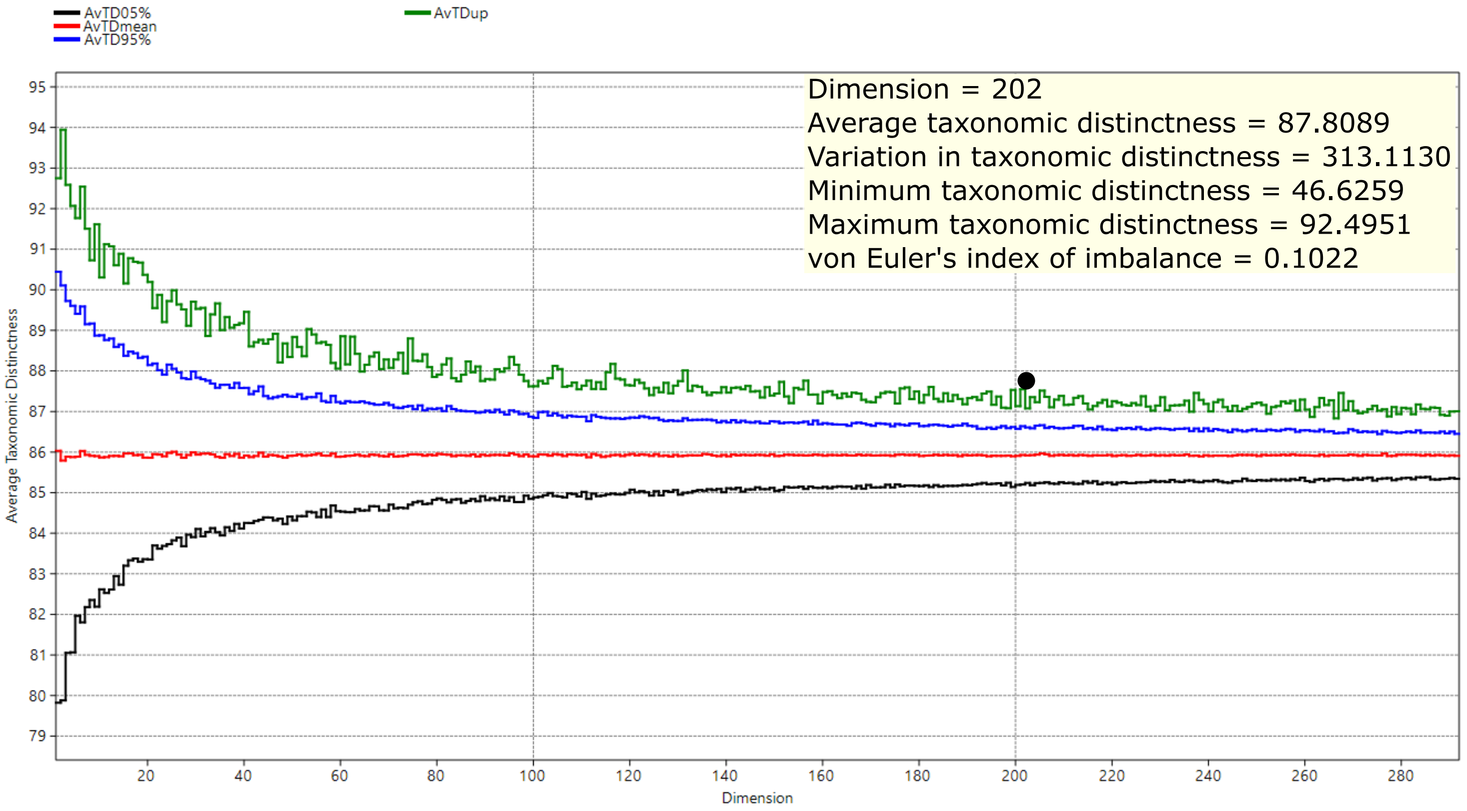
AvTD05%
AvTDmean
AvTD95%
AvTDup
Dimension = 202
Average taxonomic distinctness = 87.8089
Variation in taxonomic distinctness = 313.1130
Minimum taxonomic distinctness = 46.6259
Maximum taxonomic distinctness = 92.4951
von Euler's index of imbalance = 0.1022
Average Taxonomic Distinctness
Dimension

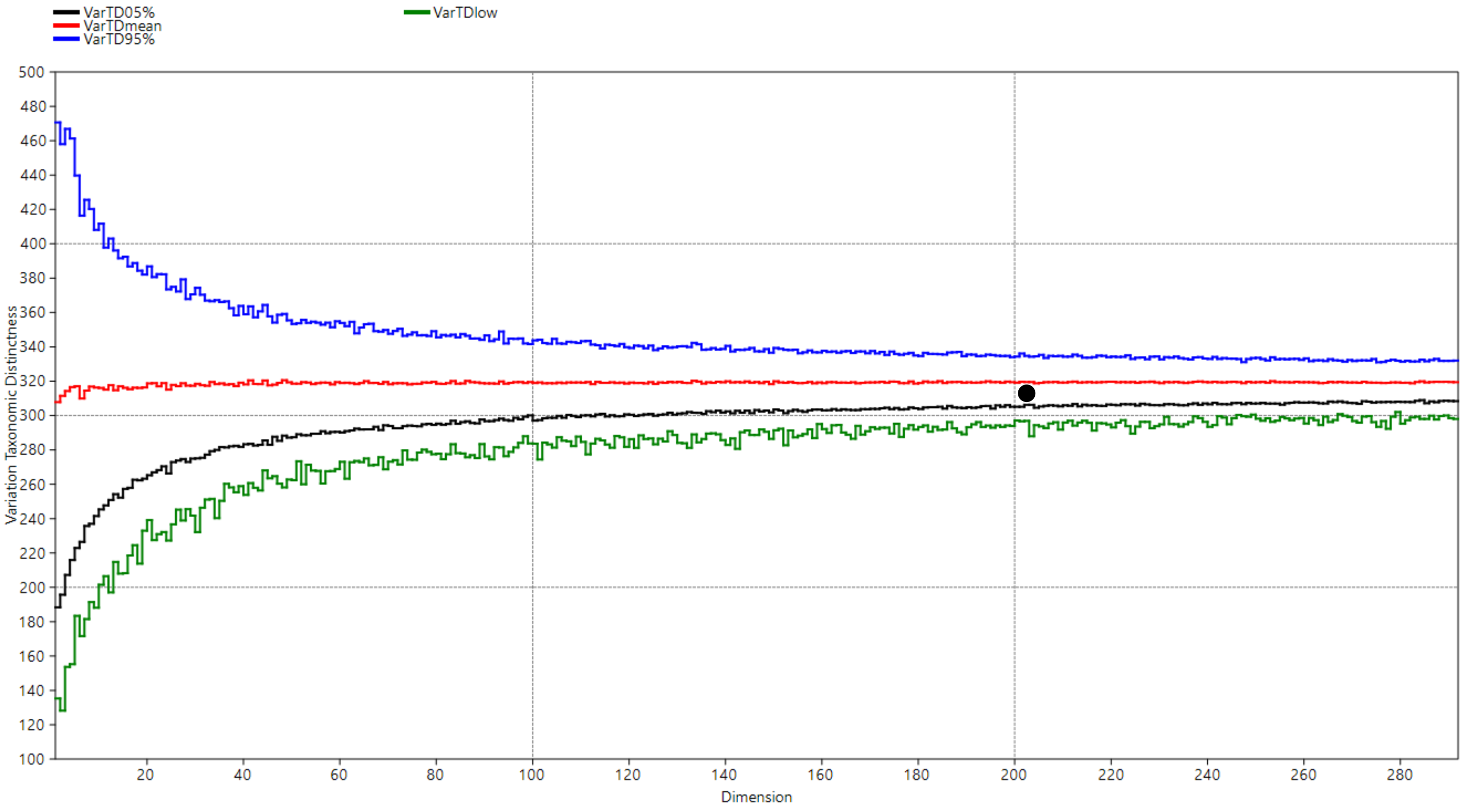
VarTD05%
VarTDmean
VarTD95%
VarTDlow
Variation Taxonomic Distinctness
Dimension

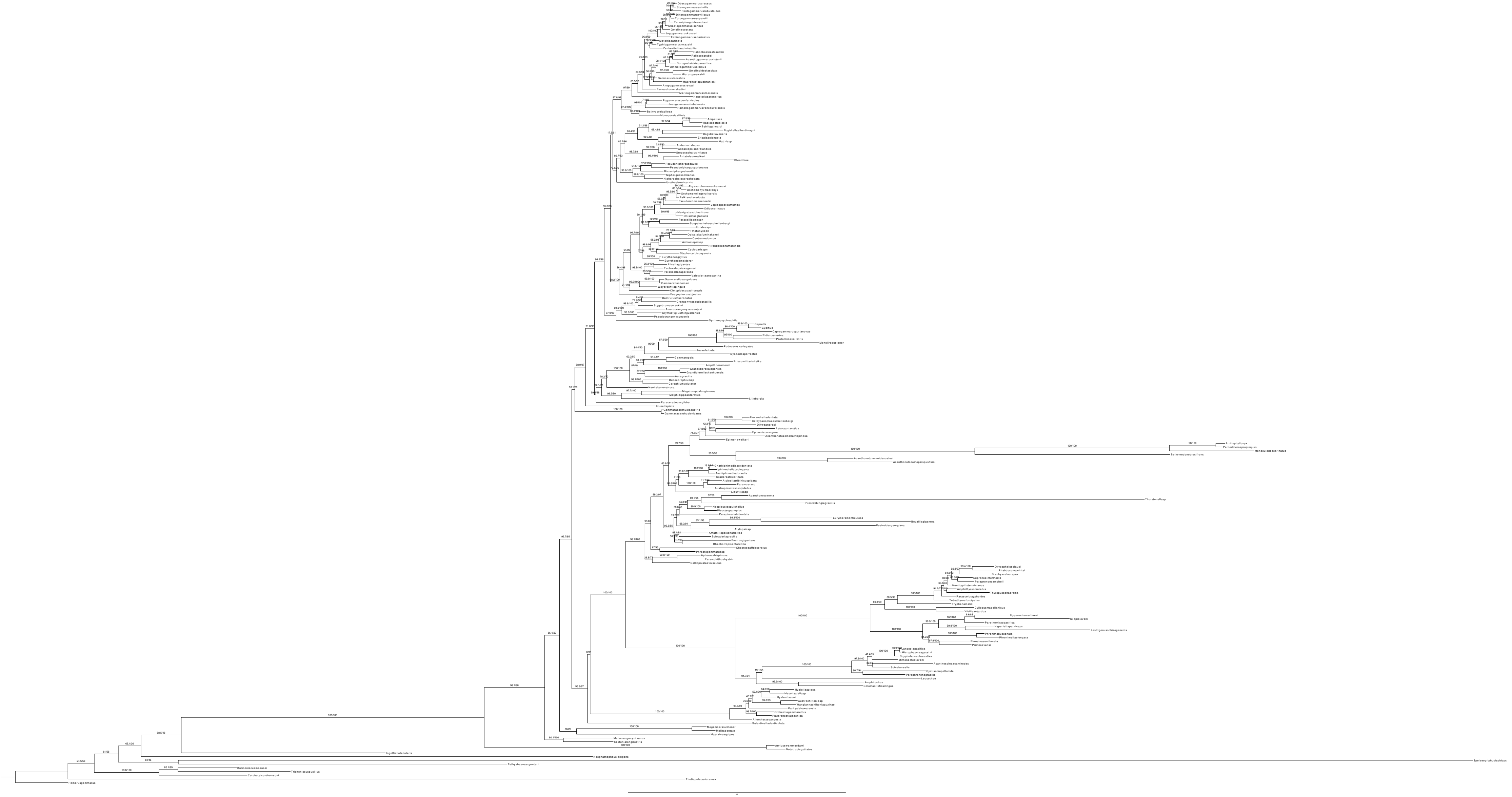

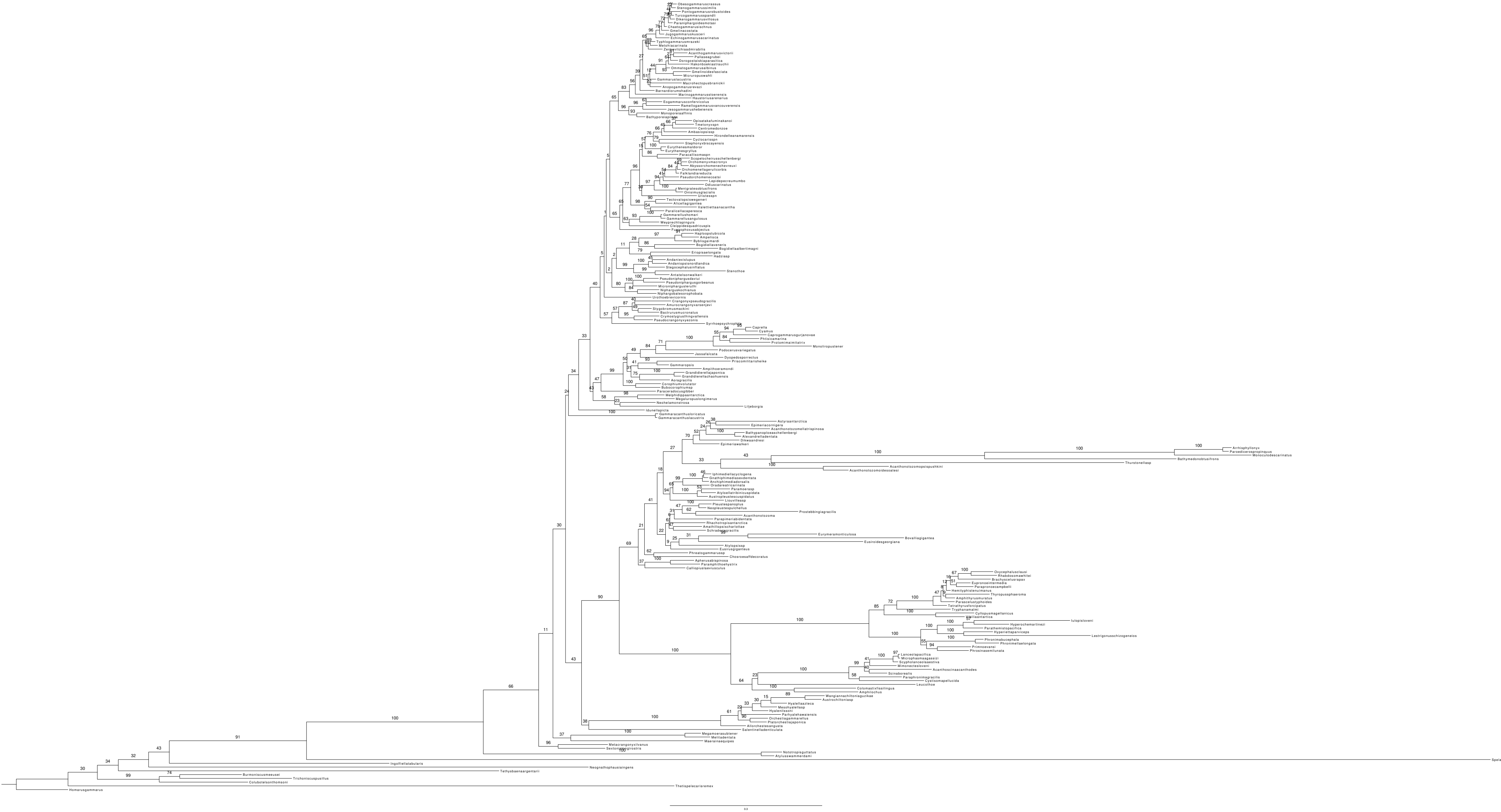
Spelaeogriphuslepidops
Ingolfiellatabularis
Neognathophausiaingens
Tethysbaenaargentarii
Thetispelecarisremex
Homarusgammarus

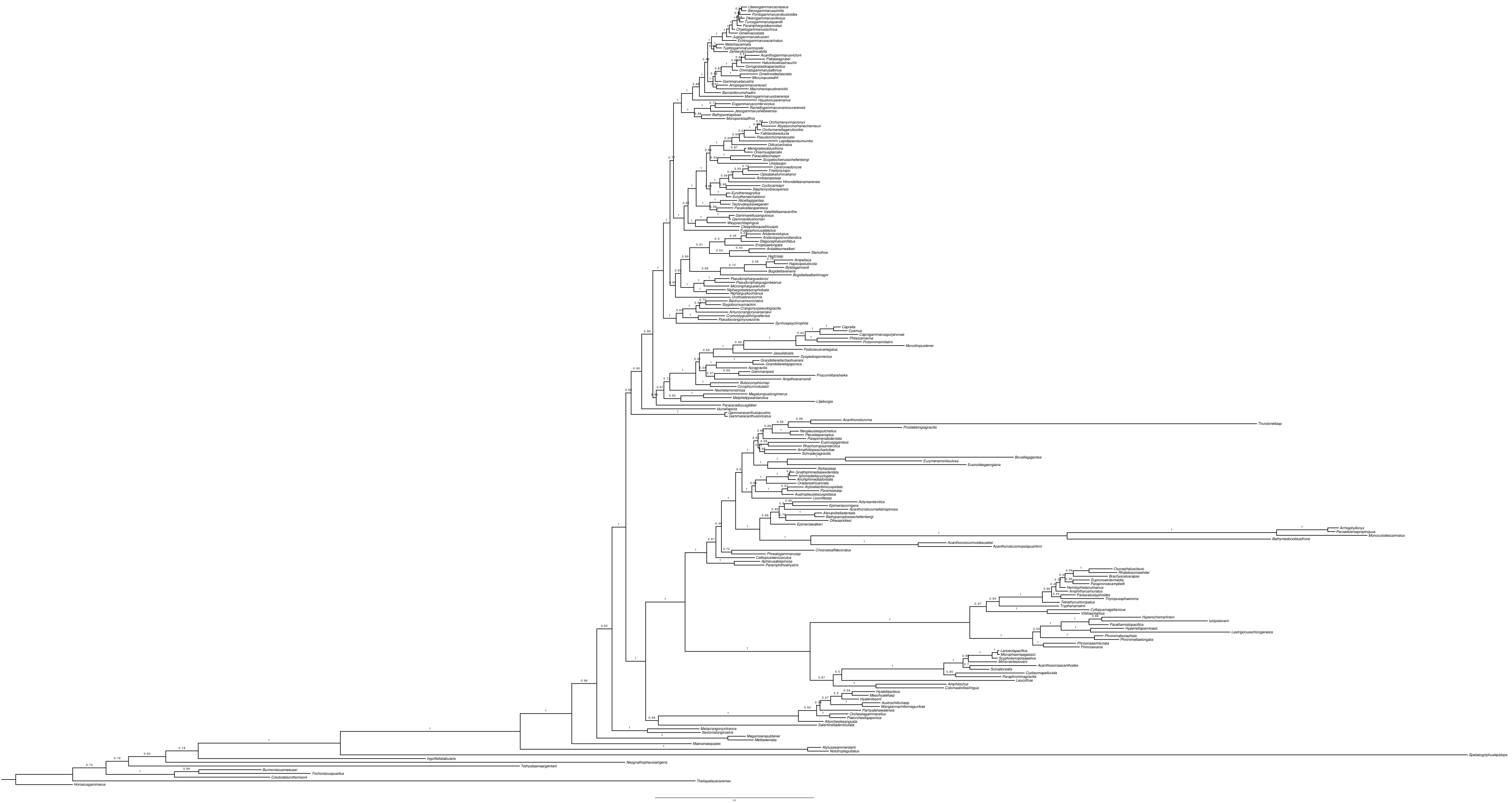

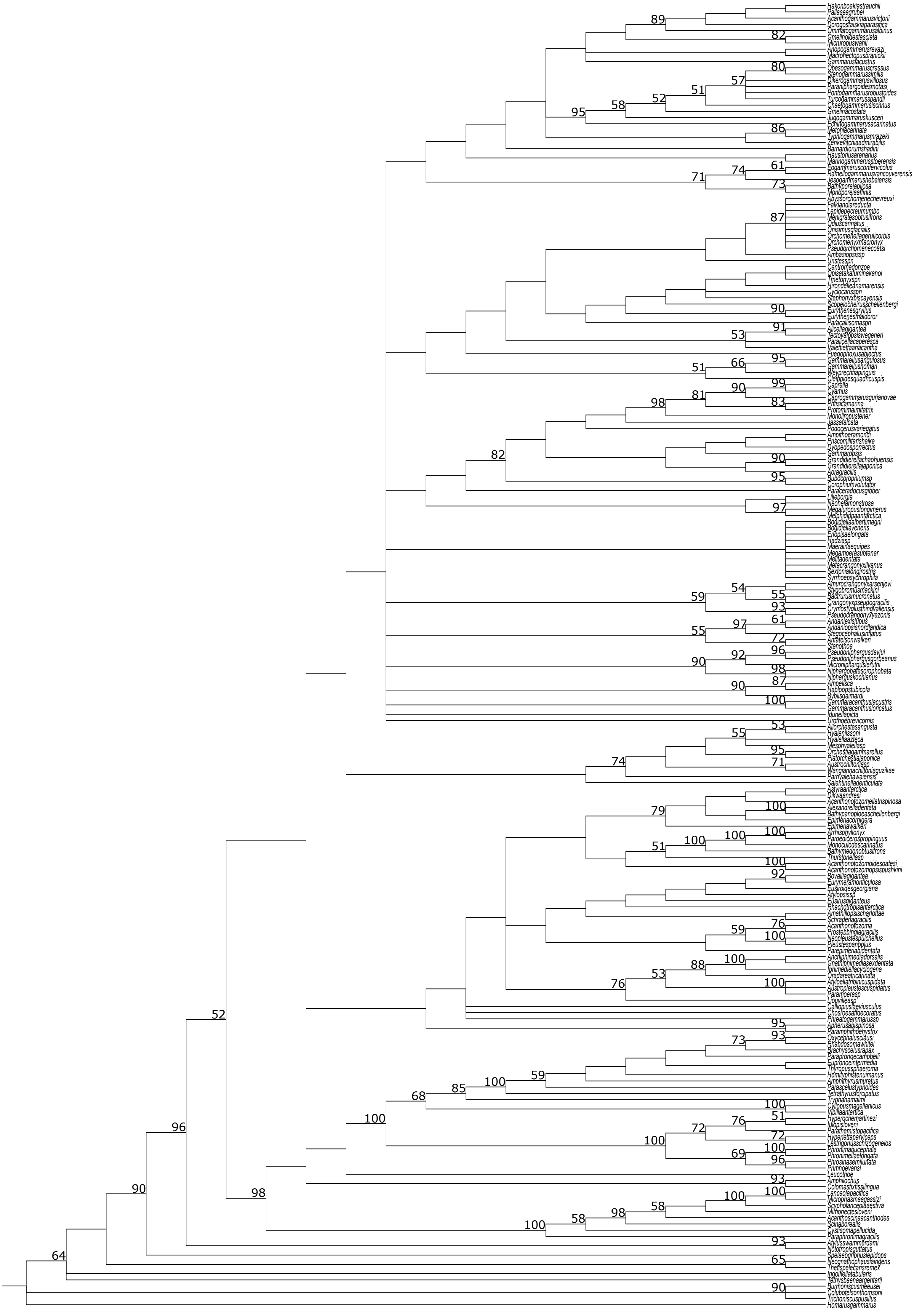
Hakonboekiastrauchii
Pallaseagrubei
Acanthogammarusvictorii
Dorogostaiskiaparasitica
Ommatogammarusalbinus
Gmelinoidesfasciata
Micruropuswahli
Anopogammarusrevazi
Macrohectopusbranickii
Gammaruslacustris
Obesogammaruscrassus
Stenogammarussimilis
Dikerogammarusvillosus
Paraniphargoidesmotasi
Pontogammarusrobustoides
Turcogammarusspandli
Chaetogammarusischnus
Gmelinacostata
Jugogammaruskusceri
Echinogammarusacarinatus
Metohiacarinata
Typhlogammarusmrazeki
Zenkevitchiaadmirabilis
Barnardiorumshadini
Haustoriusarenarius
Marinogammarusstoerensis
Eogammarusconfervicolus
Ramellogammarusvancouverensis
Jesogammarushebeiensis
Bathyporeiapilosa
Monoporeiaaffinis
Abyssorchomenechevreuxi
Falklandiareducta
Lepidepecreumumbo
Menigratesobtusifrons
Qdiuscarinatus
Onisimusglacialis
Orchomenellagerulicorbis
Orchomenyxmacronyx
Pseudorchomenecoatsi
Ambasiopsissp
Uristessp
Centromedonzoe
Opisatakafuminakanoi
Tmetonyxsp
Hirondelleanamarensis
Cyclocarissp
Stephonyxbiscayensis
Scopelocheirusschellenbergi
Eurythenesgryllus
Eurythenesmaldoror
Paracallisomasp
Alicellagigantea
Tectovalopsiswegeneri
Paralicellacaperesca
Valettiettaanacantha
Fuegophoxusabjectus
Gammarellusangulosus
Gammarellushomari
Weyprechtiapinguis
Cleippidesquadricuspis
Caprella
Cyamus
Caprogammarusgurjanovae
Phtisicamarina
Protomimaimitatrix
Monoliropustener
Jassafalcata
Podocerusvariegatus
Ampithoeramondi
Priscomilitarisheike
Dyopedosporrectus
Gammaropsis
Grandidierellachaohuensis
Grandidierellajaponica
Aoragracilis
Bubocorophiumsp
Corophiumvolutator
Paraceradocusgibber
Lilljeborgia
Neohelamonstrosa
Megaluropuslongimerus
Melphidippaantarctica
Bogidiellaalbertimagni
Bogidiellaveneris
Eriopisaelongata
Hadziasp
Maerainaequipes
Megamoerasubtener
Melitadentata
Metacrangonyxilvanus
Sextonialongirostris
Syrrhoepsychrophila
Amurocrangonyxarsenjevi
Stygobromusmackini
Bactrurusmucronatus
Crangonyxpseudogracilis
Crymostygiusthingvallensis
Pseudocrangonyxyezonis
Andaniexislupus
Andaniopsisnordlandica
Stegocephalusinflatus
Antatelsonwalkeri
Stenothoe
Pseudoniphargusdaviui
Pseudoniphargusgorbeanus
Microniphargusleruthi
Niphargobatesorophobata
Niphargus kochianus
Ampelisca
Haploopstubicola
Byblisgaimardi
Gammaracanthuslacustris
Gammaracanthusloricatus
Idunellapicta
Urothoebrevicornis
Allorchestesangusta
Hyalenilssoni
Hyalellaazteca
Mesohyalellasp
Orchestiagammarellus
Platorchestiajaponica
Austrochiltoniasp
Wangiannachiltoniaguzikae
Parhyalehawaiensis
Salentinelladenticulata
Astyraantarctica
Dikwaandresi
Acanthonotozomellatrispinosa
Alexandrelladentata
Bathypanoploeaschellenbergi
Epimeriacornigera
Epimeriawalkeri
Arrhisphyllonyx
Paroediceropspropinquus
Monoculodescarinatus
Bathymedonobtusifrons
Thurstonellasp
Acanthonotozomoidesoatesi
Acanthonotozomopsispushkini
Bovalliagigantea
Eurymeramonticulosa
Eusiroidesgeorgiana
Atylopsissp
Eusirusgiganteus
Rhachotropisantarctica
Amathillopsischarlottae
Schraderiagracilis
Acanthonotozoma
Prostebbingiagracilis
Neopleustespulchellus
Pleustespanoplus
Parepimeriabidentata
Anchiphimediadorsalis
Gnathiphimediasexdentata
Iphimediellacyclogena
Oradareatricarinata
Atyloellatricuspidata
Austropleustescuspidatus
Paramperasp
Liouvilleasp
Calliopiuslaeviusculus
Chosroesaffdecoratus
Phreatogammarussp
Apherusabispinosa
Paramphithoehystrix
Oxycephalusclausi
Rhabdosomawhitei
Brachyscelusrapax
Parapronoecampbelli
Eupronoeintermedia
Thyropussphaeroma
Hemityphistenuimanus
Amphithyrusmuratus
Parascelustyphoides
Tetrathyrusforcipatus
Tryphanamalmi
Cyllopusmagellanicus
Vibiliaantarctica
Hyperochemartinezi
Iulopisloveni
Parathemistopacifica
Hyperiettaparviceps
Lestrigonusschizogeneios
Phronimabucephala
Phronimellaelongata
Phrosinasemilunata
Primnoevansi
Leucothoe
Amphilochus
Colomastixfissilingua
Lanceolapacifica
Microphasmaagassizi
Scypholanceolaaestiva
Mimonectesloveni
Acanthoscinaacanthodes
Scinaborealis
Cystisomapellucida
Paraphronimagracilis
Atylusswammerdami
Nototropisguttatus
Spelaeogriphuslepidops
Neognathophausiaingens
Thetispelecarisremex
Ingolfiellatabularis
Tethysbaenaargentarii
Burrioniscusmeeusei
Colubotelsonthomsoni
Trichoniscuspusillus
Homarusgammarus

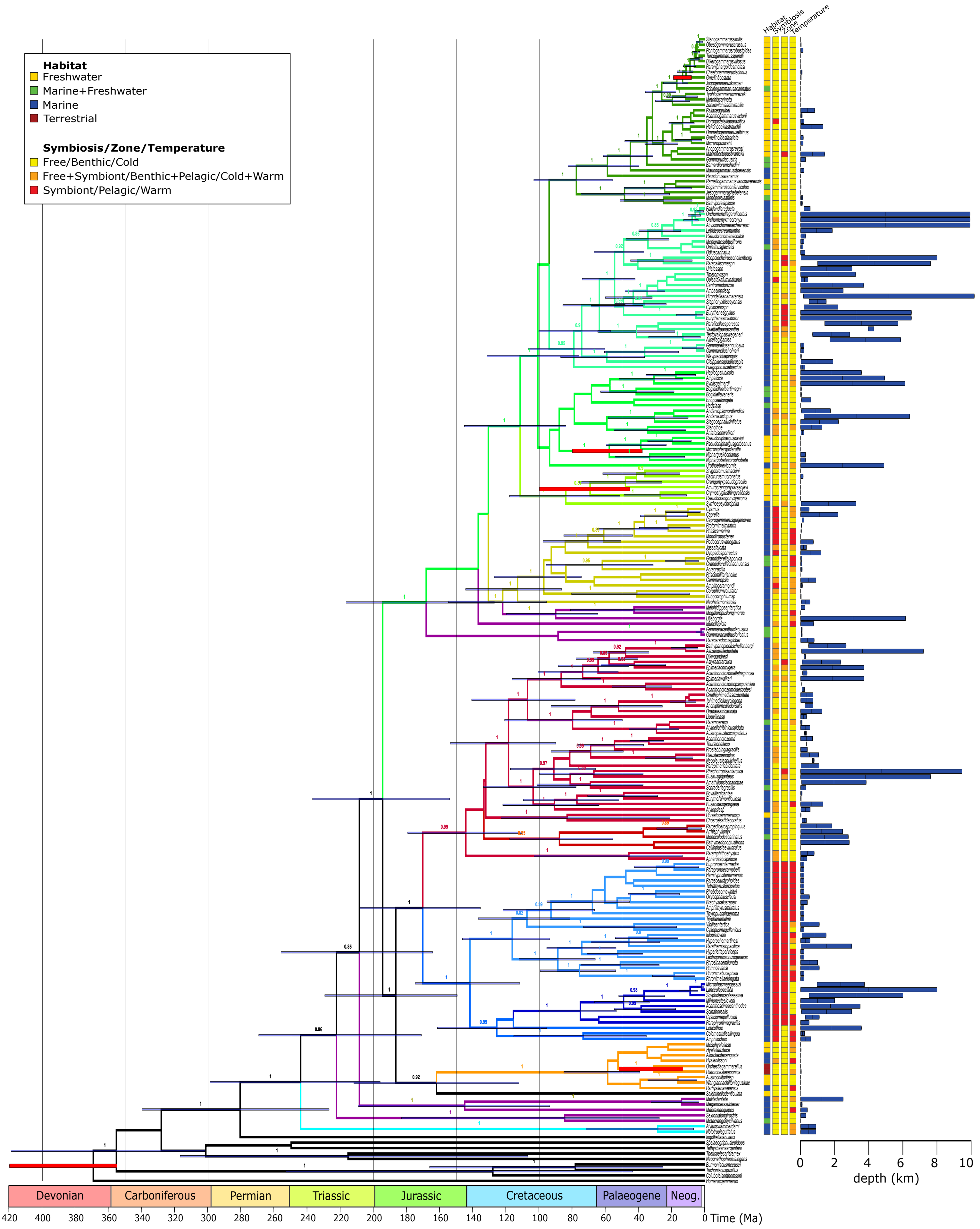
Habitat
Freshwater
Marine+Freshwater
Marine
Terrestrial
Symbiosis/Zone/Temperature
Free/Benthic/Cold
Free+Symbiont/Benthic+Pelagic/Cold+Warm
Symbiont/Pelagic/Warm
Habitat
Symbiosis
Zone
Temperature
Devonian
Carboniferous
Permian
Triassic
Jurassic
Cretaceous
Palaeogene
Neog.
420 400 380 360 340 320 300 280 260 240 220 200 180 160 140 120 100 80 60 40 20 0
Time (Ma)
0 2 4 6 8 10
depth (km)

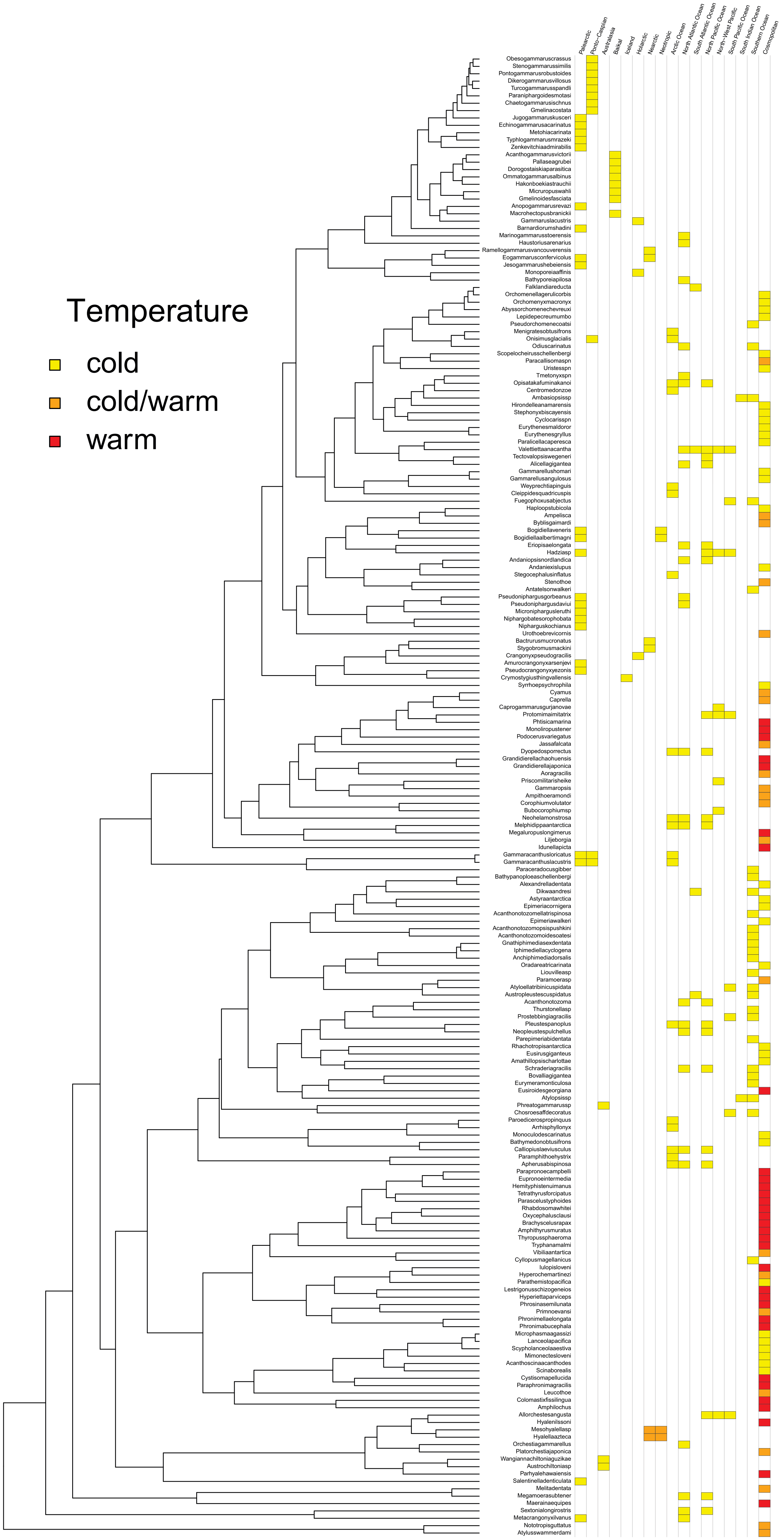

Temperature
cold
cold/warm
warm
Palearctic
Ponto-Caspian
Australasia
Baikal
Iceland
Holarctic
Nearctic
Neotropic
Arctic Ocean
North Atlantic Ocean
South Atlantic Ocean
North Pacific Ocean
North-West Pacific
South Pacific Ocean
South Indian Ocean
Southern Ocean
Cosmopolitan
Obesogammaruscrassus
Stenogammarussimilis
Pontogammarusrobustoides
Dikerogammarusvillosus
Turcogammarusspandli
Paraniphargoidesmotasi
Chaetogammarusischnus
Gmelinacostata
Jugogammaruskusceri
Echinogammarusacarinatus
Metohiacarinata
Typhlogammarusmrazeki
Zenkevitchiaadmirabilis
Acanthogammarusvictorii
Pallaseagrubei
Dorogostaiskiaparasitica
Ommatogammarusalbinus
Hakonboekiastrauchii
Micruropuswahli
Gmelinoidesfasciata
Anopogammarusrevazi
Macrohectopusbranickii
Gammaruslacustris
Barnardiorumshadini
Marinogammarusstoerensis
Haustoriusarenarius
Ramellogammarusvancouverensis
Eogammarusconfervicolus
Jesogammarushebeiensis
Monoporeiaaffinis
Bathyporeiapilosa
Falklandiareducta
Orchomenellagerulicorbis
Orchomenyxmacronyx
Abyssorchomenechevreuxi
Lepidepecreumumbo
Pseudorchomenecoatsi
Menigratesobtusifrons
Onisimusglacialis
Odiuscarinatus
Scopelocheirusschellenbergi
Paracallisomaspn
Uristesspn
Tmetonyxspn
Opisatakafuminakanoi
Centromedonzoe
Ambasiopsissp
Hirondelleanamarensis
Stephonyxbiscayensis
Cyclocarisspn
Eurythenesmaldoror
Eurythenesgryllus
Paralicellacaperesca
Valettiettaanacantha
Tectovalopsiswegeneri
Alicellagigantea
Gammarellushomari
Gammarellusangulosus
Weyprechtiapinguis
Cleippidesquadricuspis
Fuegophoxusabjectus
Haploopstubicola
Ampelisca
Byblisgaimardi
Bogidiellaveneris
Bogidiellaalbertimagni
Eriopisaelongata
Hadziasp
Andaniopsisnordlandica
Andaniexislupus
Stegocephalusinflatus
Stenothoe
Antatelsonwalkeri
Pseudoniphargusgorbeanus
Pseudoniphargusdaviui
Microniphargusleruthi
Niphargobatesorophobata
Niphargiskochianus
Urothoebrevicornis
Bactrurusmucronatus
Stygobromusmackini
Crangonyxpseudogracilis
Amurocrangonyxarsenjevi
Pseudocrangonyxyezonis
Crymostygiusthingvallensis
Syrrhoepsychrophila
Cyamus
Caprella
Caprogammarusgurjanovae
Protomimaimitatrix
Phtisicamarina
Monoliropustener
Podocerusvariegatus
Jassafalcata
Dyopedosporrectus
Grandidierellachaohuensis
Grandidierellajaponica
Aoragracilis
Priscomilitarisheike
Gammaropsis
Ampithoeramondi
Corophiumvolutator
Bubocorophiumsp
Neohelamonstrosa
Melphidippaantarctica
Megaluropuslongimerus
Liljeborgia
Idunellapicta
Gammaracanthusloricatus
Gammaracanthuslacustris
Paraceradocusgibber
Bathypanoploeaschellenbergi
Alexandrelladentata
Dikwaandresi
Astyraantarctica
Epimeriacornigera
Acanthonotozomellatrispinosa
Epimeriawalkeri
Acanthonotozomopsispushkini
Acanthonotozomoidesoatesi
Gnathiphimediasexdentata
Iphimediellacyclogena
Anchiphimediadorsalis
Oradareatricarinata
Liouvilleasp
Paramoerasp
Atyloellatribinicuspidata
Austropleustescuspidatus
Acanthonotozoma
Thurstonellasp
Prostebbingiagracilis
Pleustespanoplus
Neopleustespulchellus
Parepimeriabidentata
Rhachotropisantarctica
Eusirusgiganteus
Amathillopsischarlottae
Schraderiagracilis
Bovalliagigantea
Eurymeramonticulosa
Eusiroidesgeorgiana
Atylopsissp
Phreatogammarussp
Chosroesaffdecoratus
Paroediceropropinquus
Arrhisphyllonyx
Monoculodescarinatus
Bathymedonobtusifrons
Calliopiuslaeviusculus
Paramphithoehystrix
Apherusabispinosa
Parapronoecampbelli
Eupronoeintermedia
Hemityphistenuimanus
Tetrathyrusforcipatus
Parascelustyphoides
Rhabdosomawhitei
Oxycephalusclausi
Brachyscelusrapax
Amphithyrusmuratus
Thyropussphaeroma
Tryphanamalmi
Vibiliaantartica
Cyllopusmagellanicus
Iulopisloveni
Hyperochemartinezi
Parathemistopacifica
Lestrigonusschizogeneios
Hyperiettaparviceps
Phrosinasemilunata
Primnoevansi
Phronimellaelongata
Phronimabucephala
Microphasmaagassizi
Lanceolapacifica
Scypholanceolaaestiva
Mimonectesloveni
Acanthoscinaacanthodes
Scinaborealis
Cystisomapellucida
Paraphronimagracilis
Leucothoe
Colomastixfissilingua
Amphilochus
Allorchestesangusta
Hyalenilssoni
Mesohyalellasp
Hyalellaazteca
Orchestiagammarellus
Platorchestiajaponica
Wangiannachiltoniaguzikae
Austrochiltoniasp
Parhyalehawaiensis
Salentinelladenticulata
Melitadentata
Megamoerasubtener
Maerainaequipes
Sextonialongirostris
Metacrangonyxilvanus
Nototropisguttatus
Atylusswammerdami

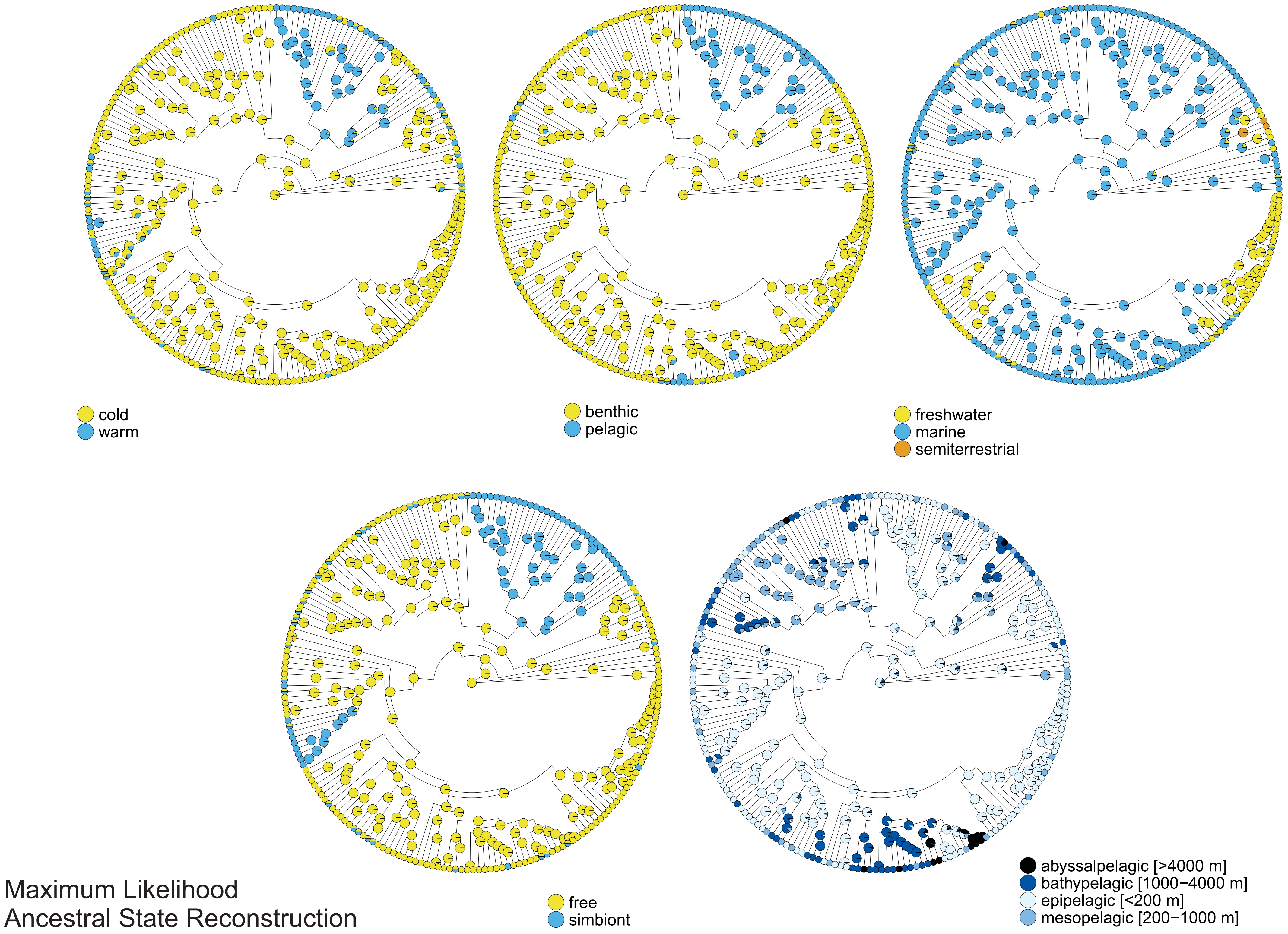
cold
warm
benthic
pelagic
freshwater
marine
semiterrestrial
free
simbiont
abyssalpelagic [>4000 m]
bathypelagic [1000−4000 m]
epipelagic [<200 m]
mesopelagic [200−1000 m]
Maximum Likelihood
Ancestral State Reconstruction
